# Ablation of a maternal *Cryptosporidium* mRNA-binding protein results in sterile sporozoites

**DOI:** 10.64898/2026.09.01.748511

**Authors:** Abigail M. Daniels, Sebastian Shaw, Christopher Nötzel, Rui Xiao, Keenan O’Dea, Daniel P. Beiting, Boris Striepen

## Abstract

Infection with *Cryptosporidium* is a leading cause of diarrheal disease and early childhood mortality. This apicomplexan parasite undergoes asexual and sexual replication within the same host and recent studies have shown an intrinsic developmental program of obligate transition to male and female gametes and sex. While factors were identified that control male fate and development, how female gene expression is orchestrated remains largely unknown. Here we use the *Cryptosporidium* Single Cell Atlas to discover an RNA binding protein (F-RBP) as one of the earliest markers of female identity. Reporter parasites engineered based on this gene allowed us to calibrate transcriptional pseudotime against the real time of female development revealing a significant window of transcriptional fate ambiguity. While F-RBP is an early transcript, the protein persists throughout female development and into the zygote. Conditional ablation of the F-RBP gene showed it to be dispensable for sex determination and early female development *in vitro*. However, the gene is essential *in vivo* and its loss results in rapid cure. Cell biological experiments link this loss to the production of sterile oocysts which release sporozoites incapable of host cell invasion. F-RBP binds transcripts highly expressed in the female gamete enriched for a YBOX primary sequence motif and forms mRNA protein complexes in late females akin to processing or P bodies. We propose F-RBP’s essential role to be in the regulation of long-term homeostasis of maternally inherited RNA required for sporozoite infectivity.

**Importance:** The parasite *Cryptosporidium* is responsible for millions of cases of severe diarrheal disease in young children and is both a consequence of, and a risk factor for, chronic malnutrition. There are no vaccines available and a single approved treatment is largely ineffective. Infection relies on ingestion of the oocyst, an environmentally stable spore, which contains an invasive form of the parasite, the sporozoite. Oocyst and sporozoites are the product of parasite sex and the fusion of female and male *Cryptosporidium* parasites. Here we report the critical role of an RNA-binding protein in female mRNA homeostasis. Maternal transcripts orchestrate the complex process of sexual replication and oocyst formation. Loss of the protein results in sterile spores that fail to initiate the next round of the lifecycle, and conditional ablation using a small molecule trigger quickly cures infected animals demonstrating the essential role of parasite sex in continued infection.

## Introduction

Sexual reproduction is a hallmark of eukaryotic life. In most systems, two distinct and often differently sized cells contribute to this process: the male cell (sperm) and the female cell (egg). The sperm’s primary role is to deliver its genetic material. The egg is responsible for enabling male recognition to allow fertilization, prevention of polyspermy, early embryogenesis, and often the storage and provision of the metabolic reserves to fuel these activities. Egg cells harbor a wide array of biological materials and mRNAs are an important part of this cargo as they encode proteins needed throughout egg development and early embryogenesis(1, 2). Carrying these mRNAs allows the fertilized egg to execute a series of rapid transitions without the need for transcription. Multicellular and unicellular eukaryotes alike use RNA binding proteins to ensure that these messages are stored, protected, and transcribed exactly when needed in development(1–4).

Apicomplexan parasites similarly rely on sexual reproduction, and this is of particular importance to *Cryptosporidium*. This parasite goes through a defined series of asexual replication cycles that culminate in the formation of the sperm and egg equivalent, the male and female gametes, respectively(5). The male gamete fertilizes the female gamete, giving rise to the oocyst, a spore-like stage which can transmit the infection to a new host, or remain within the original host to continue the infection.

Akin to other female reproductive cells, the *Cryptosporidium* female gamete prepares for different events at distinct developmental stages. First, this cell needs to commit to, and initiate, the female genetic program. During this early stage of female development, the parasite grows significantly and stockpiles metabolites, proteins, and mRNA to prepare for fertilization and sporulation. *Cryptosporidium* is a haplont and will enter meiosis immediately after pronuclear fusion, and the female gamete carries the required machinery(6–10). Maternal mRNA is also required for the assembly of the crystalloid body(7), the synthesis of storage polymers(11), sporozoite assembly(12), and the biogenesis of the oocyst wall(13). The oocyst wall is composed of glycans, specialized lipids and about 40 proteins including the *Cryptosporidium* Oocyst Wall Proteins (COWPs), which are expressed during female development in two distinct waves (6, 13–16). Given suitable conditions, the oocyst can lay dormant for months(17), and upon ingestion, release motile sporozoites which invade the intestinal epithelium to colonize a new host.

Sex is of particular importance for *Cryptosporidium* as it sustains continued infection(6, 7, 9–11, 18, 19). In standard cell culture gametes are formed but fertilization does not occur and no growth is observed past two days(11). Organoid model systems allow for oocyst production, though not yet at the levels observed in mice(19). This is the consequence of a hard-wired lifecycle and obligate switch towards sex (5, 8, 18). Remarkably, preventing parasite sex will rapidly halt the infection in the same host (6).

Apetala 2 (AP2) transcription factors mediate developmental changes in a variety of apicomplexan parasites and are important regulators of gene expression, especially during sexual development(20–22). The female gamete expresses three of these transcription factors(6). One has been implicated in development of the sporozoite specific organelle, the crystalloid body(7), while another has been linked to female gamete development *in vitro*(23). However, all three of these transcription factors are transcribed at a similar late point late in development providing only limited time resolution(6).

Here, we characterize a female specific RNA binding protein, F-RBP. F-RBP preferentially binds mRNAs that are upregulated in female development and the absence of F-RBP results in female sterility and sporozoites incapable of host cell infection.

## Results

### Female identity is transcriptionally evident only at the half point of gamete development

We mined the *Cryptosporidium* female exclusive single cell transcriptome(6) for potential regulators of female gene expression, focusing on those genes expressed very early, and exclusively, during female development(12). We identified two genes: cgd4_130, encoding a putative RNA Recognition Motif (RRM) and K Homology (KH) RNA binding protein and cgd4_1360, a Zinc Finger domain containing protein (referred to as F-RBP and F-ZF, respectively, Figure 1A). Single cell RNA profiling has provided significant insight into the relative sequence of gene expression in *Cryptosporidium,* but we do not understand how the pseudotime derived from these analyses translates into the actual time of parasite development. To sync these two clocks and to monitor development over time, we constructed a fluorescent reporter strain that combined the *f-rbp* promoter to control expression of the green fluorescent protein mNeon and the promoter of the previously characterized *cowp1*(11) to control expression of the red fluorescent protein tdTomato (Figure 1B). Stable transgenic parasites were selected with paromomycin, and genome modification was validated by PCR mapping of genomic DNA isolated from oocysts purified from the feces of infected mice (Supplemental Figure 1A)(24).

**Figure 1:**
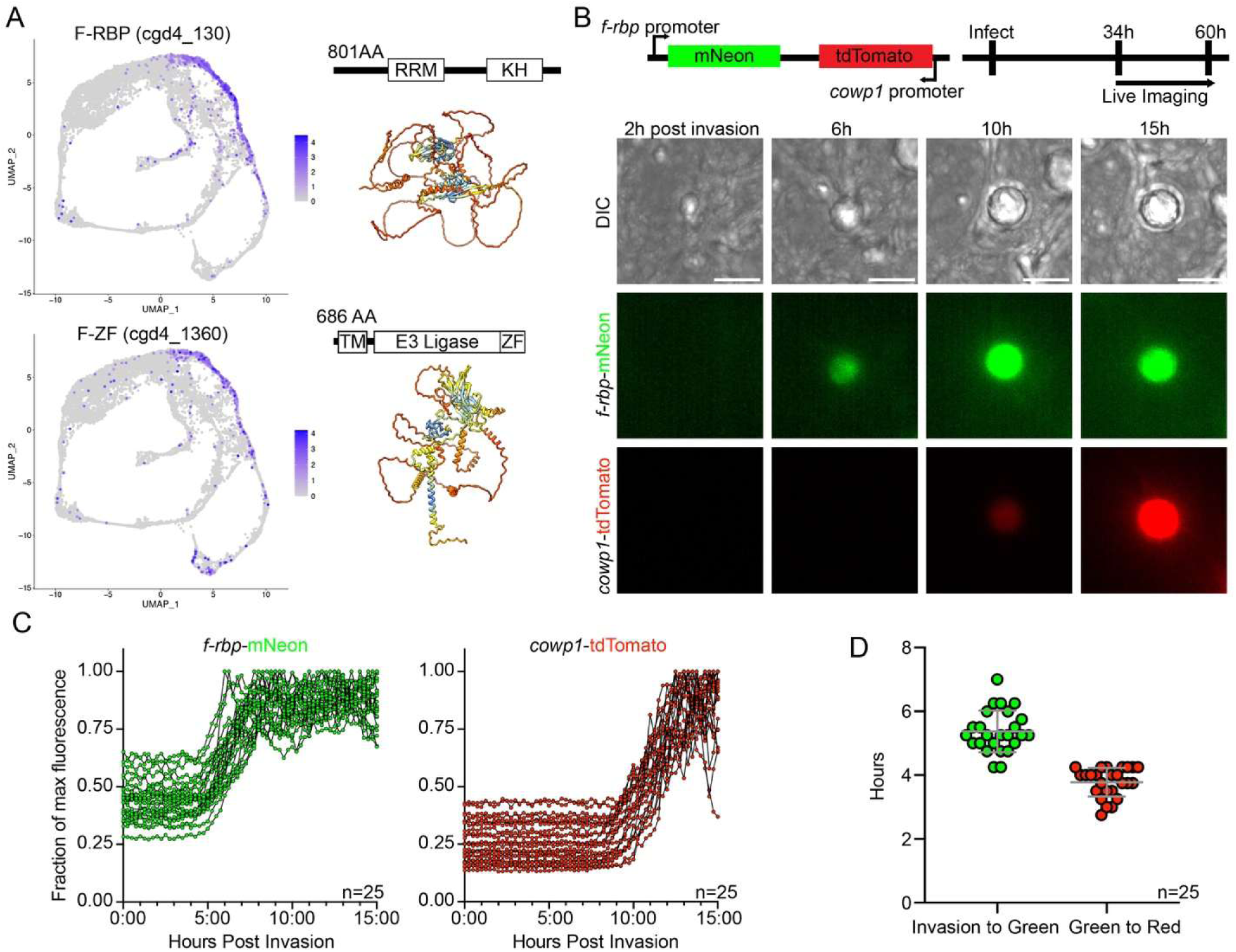
F-RBP expression defines early female gamete development. **A.** Transcript expression mapped onto the Single Cell Atlas(6), domain and Alphafold structure prediction of F-RBP (top) and F-ZF (bottom) **B.** Construct and experimental design for female reporter live imaging assay (top). Representative fluorescence and differential interference contrast (DIC) images of a female gamete over time. Scale bar 5µm. **C.** Fluorescence intensity plotted as fraction of maximum fluorescence over time for *f-rbp-*mNeon and *cowp1-*tdTomato. Each continuous line represents a single tracked parasite. n=25. **D.** Time measured between indicated event for each of the 25 gametes. First detection was assigned to the time when fluorescence intensity exceeded the mean background by 5 times the SD. Mean and SD indicated in gray. n=25.

Transgenic parasites were used to infect the human adenocarcinoma cell line HCT-8 seeded into glass bottom chamber slides and subjected to time lapse imaging after 34 h of growth (see Methods for technical detail). At this point parasites egress from their final round of asexual replication, followed by reinvasion and differentiation into gametes(6, 18). We detected no fluorescence in either channel for invading merozoites (Figure 1B&C, Movie S1). When tracking the resulting intracellular stages, mNeon driven by the *f-rbp* promoter was detectable first, followed later by *cowp1*-driven tdTomato. We recorded the development of a total of 25 female gametes. Parasites were segmented in each frame using the DIC images as a mask, and we measured and plotted the fluorescence intensity for both reporters over time. We found that the timing of reporter expression was remarkably consistent (Figure 1C). After the parasites invade, it takes 5 h and 20 min (SD ± 40 min) for the *f-rbp* promoter to turn on, marking them as female. It then takes another 3 hours and 45 minutes (± 30 min) to detect *cowp*1 driven fluorescence (Figure 1D). We thus note a substantial phase in which the lifecycle stage remains transcriptionally undifferentiated. After this period, gametogenesis proceeds through precisely timed waves of expression of female genes where we time the *cowp1* expression to 9h of the roughly 12h gamete development.

### F-RBP and F-ZF proteins are translated early in female development

To assess the level of F-RBP and F-ZF proteins over the course of parasite development, we introduced a DNA cassette encoding a 3xHA epitope tag to modify the C terminus of each protein into the native gene locus (Supplemental Figure 1B&C). HCT-8 cultures were infected with the resulting transgenic parasites and processed for immunofluorescence microscopy at different time points. No labeling was observed for F-RBP-HA or F-ZF-HA after 24h of infection, in accordance with our transcriptomic and reporter results (Figure 2A-C). F-RBP-HA was readily observed in cultures starting at 42h post infection, accounting for approximately 30% of all parasites detected by counterstain with *Vicia villosa* lectin (VVL, Figure 2A & C). This proportion increased to 50-60% of parasites at 48h post infection and remained so at 72h, consistent with the proportion of female parasites previously reported for these timepoints(11). F-ZF-HA was detected in a smaller fraction of parasites at 42 hours (0-15%), increasing to ∼30% of the population at 48 and 50% 72 hours post infection (Figure 2B & C). For both lines, HA+ parasites were observed to have a single nucleus, typical for the female gamete (11, 18) and both proteins were found consistently in parasites positive for the female marker DMC1(7, 8). DMC1 is expressed later in female development, and we therefore do detect F-RBP-HA+ parasites that are not yet expressing DMC1 (Supplemental Figure 2A). All DMC1+ parasites also stained F- RBP-HA+ (Supplemental Figure 2A).

**Figure 2:**
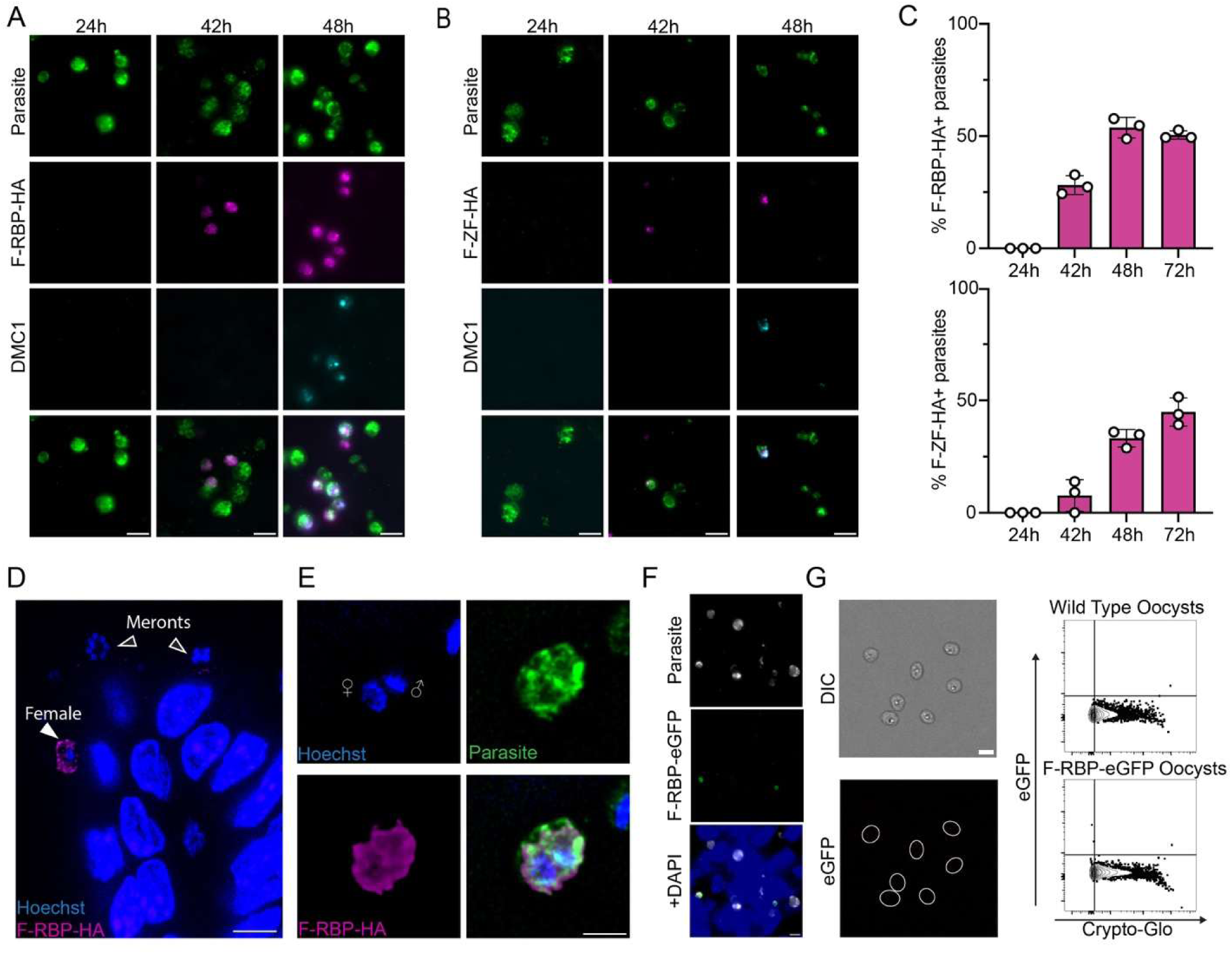
F-RBP protein is detectable from the early gamete to the zygote but absent from the oocyst. Immunofluorescence time course of **A.** F-RBP-HA and **B.** F-ZF-HA in HCT-8 tissue culture. Scale bar 5µm. Representative images from 3 independent experiments. **C.** Quantification of HA positive parasites for F-RBP-HA (top) and F-ZF (bottom). 3,438 parasites counted for F-RBP quantification and 3,920 parasites counted for F-ZF quantification. For both, 3 independent experiments with 3 coverslips per experiment were counted. Data from one representative experiment shown. **D.** F-RBP-HA staining in tissue section from infected mouse intestine. Different parasite stages are highlighted. Scale bar 5µm. **E**. Detail from tissue section showing a zygote (recognized by the presence of both a female (large) and male (small bullet-shaped) nucleus. Note that zygote stained for F-RBP-HA. Scale bar 2µm. **F.** F-RBP-eGFP parasites in HCT-8 cells. Scale bar 5µm. Note female parasites showing eGFP fluorescence. **G.** F-RBP-eGFP oocysts harvested from infected mice by microscopy (left) or flow cytometry (right). Flow cytometry is representative of 2 independent experiments. F-RBP-eGFP oocysts show no fluorescence over control (wild type). Scale bar 5µm.

### F-RBP is present in zygotes, but is absent in oocysts

While female gametes form in HCT-8 cultures, they are not fertilized, precluding the observation of subsequent development(11). To investigate F-RBP expression *in vivo* we infected C57BL/6 *ifny ^-/-^* mice, and at peak infection sacrificed the animals, resected the ileum, and performed immunofluorescence on histological sections. We noted the presence of F-RBP in female gametes, identified by their single, large nucleus (Figure 2D). We also detected F-RBP in zygotes, which carry both a large female and a smaller male nucleus, demonstrating that this protein persists from early gametogenesis to fertilization and maternal-to-zygote transition (Figure 2E). We next sought to determine the fate of F-RBP in oocysts. This is technically complicated by the oocyst wall which is not readily breached by detergent permeabilization leading to inconsistent antibody access and staining. To overcome this, we engineered a parasite strain in which F-RBP was translationally fused to eGFP, thus allowing us to use endogenous fluorescence to monitor the presence of F-RBP protein (Supplemental Figure 1D).

While these parasites showed bright fluorescence in female stages at 48h *in vitro* (Figure 2F), no fluorescence was detected in oocysts by either microscopy or flow cytometry (Figure 2G). We conclude that F-RBP is expressed early in female development, maintained through gametogenesis to fertilization, but is degraded by the time the oocyst has matured.

### Genetic ablation of F-RBP and F-ZF

To assess the importance of F-RBP or F-ZF for female development and parasite lifecycle completion, we attempted to ablate each gene by homologous recombination. We readily achieved this for F-ZF, and oocysts were abundantly shed from mice infected with validated ΔF- ZF parasites indicating that this gene is dispensable for lifecycle completion (Supplemental Figure 3A&B). In contrast, parasites were not recovered in three separate attempts to ablate the C terminal portion of F-RBP, which included the KH domain (Supplemental Figure 3C). Since we were able to ablate F-ZF without apparent defect, we focused on F-RBP.

To test rigorously whether F-RBP is essential, we sought to establish a conditional knockout using rapamycin inducible Cre recombinase (6, 7, 25). However, multiple attempts to insert loxP sites were unsuccessful (Supplemental Figure 1E). Introducing double-strand breaks in close proximity to the desired site of homologous recombination can enhance efficiency(26). We tested this using two guide RNAs to cut immediately up and downstream of the region to be replaced by homologous recombination. Alternatively, we used a single guide and designed a repair template for insertion instead of replacement generating a partially pseudo-diploid floxed locus (see Figure 3A for detail on both strategies). Only the second approach proved successful (Figure 3b, Supplemental Figure 1F). We engineered F-RBP-flox parasites in two *C. parvum* backgrounds, Iowa IIa Bunchgrass and the KVI IId strain(27) and then crossed them with parasites carrying a rapamycin inducible Cre recombinase yielding F-RBP cKO parasites(25). The IIa background mutant grew poorly, and we therefore focused our efforts on the mutant in the IId background. HCT-8 cultures were infected with this mutant and incubated in the presence of rapamycin or ethanol vehicle. Rapamycin treatment resulted in complete Cre-mediated excision of the floxed sequence from the chromosomal locus over 48h and as previously described, we also noted partial excision in the absence of rapamycin due to leaky Cre activity (Figure 3C)(6, 7).

**Figure 3:**
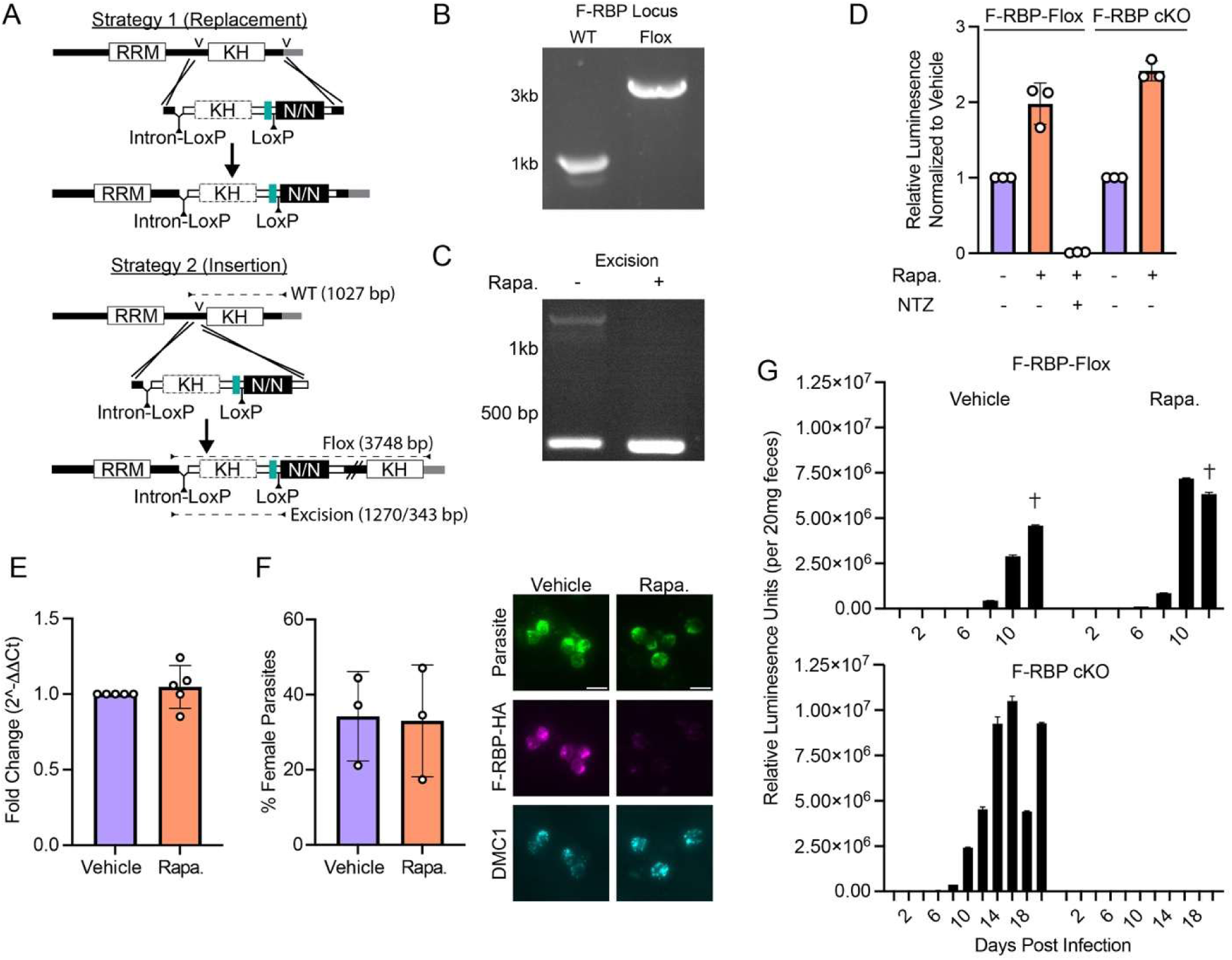
F-RBP is dispensable *in vitro*, but reduces oocyst shedding *in vivo*. **A.** Two strategies were attempted to insert loxP sites to generate a F-RBP cKO strain. In the first, the endogenous site was replaced by a floxed copy. In the second, the floxed copy was inserted into the endogenous site, pushing the remainder of the coding sequence downstream away from a start codon. Diagnostic PCR primer locations and lengths in **B** and **C** are shown in Strategy 2 by the dashed lines. gRNA locations are denoted above the gene model with arrows. **B.** Diagnostic PCR demonstrating successful modification from wild type (left) of the F-RBP locus to contain the floxed gene segment (right). **C.** Cre-mediated excision of F-RPB in parasites grown for 48h in HCT-8 cells with (right) or without Rapamycin (left). Rapamycin is abbreviated as “Rapa.” in the remainder of figure legends. We note the excision in the absence of Rapamycin, as previously reported(6, 7). **D.** Parasite growth in HCT-8 cells after 48 hours as measured by luminescence for the F-RBP-flox strain (left) and F-RBP ckO strain (right). Both were treated with either vehicle or rapamycin, which are displayed in purple and orange throughout, respectively. Raw luminescence values were normalized to vehicle treated condition within each genotype. Data shown from 3 independent experiments, each experiment performed in quintuplet. Nitazoxanide treatment served as a negative control. **E.** RT-qPCR measurement of *dmc1* transcript after 54 hours of infection with F-RBP cKO parasites in HCT-8 cells. *C. parvum gapdh* was used as the control gene. Data shown for 5 independent experiments, each experiment conducted in triplicate. **F.** Quantification of immunofluorescence of total parasites staining positive for DMC1 after 54 hours of F-RBP cKO infection in HCT-8 cells treated with vehicle or rapamycin. Note DMC1 is present even when F-RBP is absent. Representative images from 1 experiment, with 609 parasites counted across 3 coverslips per condition. Scale bar 5µm. **G.** Oocyst shedding measured from fecal luminescence from *ifny^-/-^* mice infected with F-RBP-flox parasites (top) or F-RBP cKO parasites (bottom) treated with vehicle or rapamycin from Day -2 to 10 of infection. Representative of 2 independent experiments for F-RBP cKO infection and 1 for F-RBP-flox infection. 3-5 mice per experiment. Age and sex matched within each experiment.

### F-RBP is dispensable for female gametogenesis and growth *in vitro*

We first measured the impact of F-RBP ablation on parasite growth *in vitro*. HCT-8 cells were infected with either F-RBP-flox (which lack Cre recombinase) or F-RBP cKO parasites and subjected to vehicle or rapamycin treatment. Measuring parasite burden after for 48 hours using a luciferase assay showed that rapamycin treatment did not inhibit the growth of either strain (Figure 3D). F-RBP is one of the earliest female transcripts and we therefore wondered whether it may be required for female development. HCT-8 cell cultures infected with mutant parasites were treated with vehicle or rapamycin, and we measured transcript abundance for DMC1, a meiotic protein exclusively expressed in females(7, 8). There was no significant difference between the two conditions (Figure 3E). We also assessed female development by IFA using a DMC1 antibody(7, 8) and again found no difference upon loss of F-RBP, as we readily identified parasites expressing DMC1 but lacking F-RBP-HA upon rapamycin treatment (Figure 3F).

### F-RBP ablation cures infected mice from infection

We next tested whether F-RBP may be required *in vivo*. Mice were pre-treated with either vehicle or rapamycin for 2 days and infected with F-RBP-flox or F-RBP cKO parasites, and treatment was continued for 10 days. Oocyst shedding was evaluated by measuring luciferase activity in the feces(24). Mice infected with F-RBP-flox parasites shed oocysts robustly and developed severe infection irrespective of treatment (Figure 3G). The rapamycin treated group shed more oocysts, due to immunosuppressive activity of the drug as reported previously(7). While mice infected with F-RBP cKO parasites shed oocysts throughout the duration of observation when treated with vehicle, treatment with rapamycin severely reduced oocyst shedding (Figure 3G). Nanoluciferase assays conducted using intestinal biopsy material found similarly reduced parasite burden upon rapamycin treatment in the tissue (Figure 4C). We confirmed this reduction in parasite burden by performing immunofluorescence on sections from intestinal tissue (Figure 4C). We also tested whether patently infected animals could be cured by F-RBP ablation. Mice infected with F-RBP cKO parasites were split into two groups at the peak of infection, and while vehicle treated animals remained infected, rapamycin treated animals displayed a rapid decrease in fecal luciferase signal (Supplemental Figure 4A). We conclude that F-RBP is dispensable for female development *in vitro*, but essential *in vivo* and required for sustained infection.

**Figure 4:**
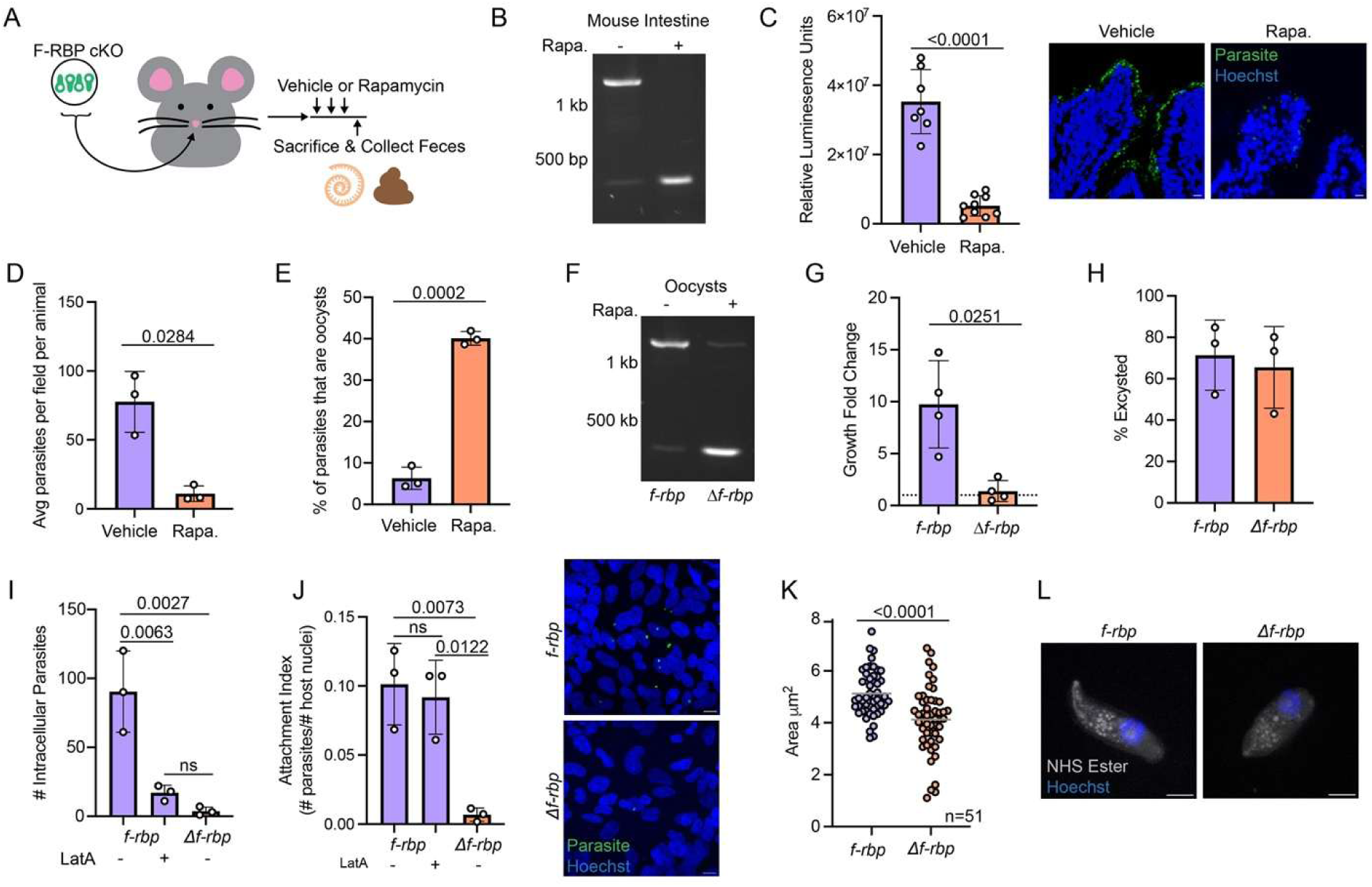
F-RBP deficient female gametes give rise to sporozoites unable to infect host cells. **A.** Experimental set up for assessing F-RBP’s absence *in vivo*. *Ifny^-/-^* mice were infected with F- RBP cKO parasites and allowed to grow for 6 days after which the animals were treated with vehicle or rapamycin daily for 72 hours. Feces were collected and the ileums were harvested the next day**. B.** Diagnostic PCR showing complete excision of F-RBP from parasites within the ileum in rapamycin treated animals (right), compared to the vehicle treated counterparts (left). Representative PCR from 2 experiments, n=7 and 9 for vehicle treated and rapamycin treated mice, respectively. **C.** Luminescence measurements from ileum punch biopsies of each vehicle or rapamycin treated animal (left). Data combined from 2 independent experiments. Same animal number as **B**. 3 punch biopsies collected per animal, and measured in triplicate. Displayed value calculated as average of each biopsy. Welch’s t-test used to determine significance. Representative images of parasite burden in intestinal tissue section (right). Scale bar 10µm. **D.** Number of parasites counted using immunofluorescence from intestinal tissue sections. The average number of parasites counted per field of view for each animal (3 per group) is displayed. 1,230 parasites counted in total. Welch’s t-test used to determine significance. **E.** Percentage of parasites counted in **D** exhibited Crypto-Glo staining consistent with oocyst stage. Welch’s t test used to determine significance. **F.** Diagnostic PCR for F-RBP excision using DNA isolated from oocysts shed from mice infected with F-RBP cKO parasites and treated with vehicle or rapamycin. Resulting parasites shed from vehicle and rapamycin treated animals referred to as *f- rbp* or Δ*f-rbp*, respectively. **G.** Growth of *f-rbp* or Δ*f-rbp* parasites after 48 hours in HCT-8 cells measured by luminescence. Fold change growth was calculated by dividing a 48h measurement by the 4h measurement per genotype. Data combined from 4 independent experiments, all measured in quintuplet. Welch’s t test used to determine significance. **H.** Percentage of *f-rbp* or Δ*f-rbp* oocysts that have excysted sporozoites. Oocyst were counted, allowed to excyst, and number of sporozoites counted to determine excystation percentage. Data combined from 3 independent experiments. 1287 sporozoites counted between 3 independent experiments. **I.** Quantification of intracellular parasites after *f-rbp* or Δ*f-rbp* infection in HCT-8 cells after 2 hours. An *f-rbp* sample was treated with Latrunculin A (LatA) to inhibit invasion. Data from one representative experiment shown of 3 for *f-rbp* or Δ*f-rbp*, and 2 experiments for *f-rbp* + LatA, with each experiment comprising of 3 coverslips. 1,627 parasites counted across all experiments. **J.** Attachment index from images collected in **I** calculated as # total parasites/# host cells (left). 19,699 host cells counted across all experiments. One way ANOVA with multiple comparisons used to determine significance for **I** and **J**. Representative images of *f-rbp* or Δ*f-rbp* infected HCT-8 cells field of view (right). Scale bar 10 µm. **K.** Measured area in µm^2^ of *f-rbp* and Δ*f-rbp* sporozoites. 51 parasites per group were measured. Welch’s t test was used to measure significance. **L.** Ultrastructure Expansion Microscopy (U-ExM) images of *f-rbp* or Δ*f-rbp* sporozoites. Note the Δ*f-rbp* sporozoites are smaller, stouter, and have a less defined apical end. Expansion factor of 4.42. Scale bar 1 µm.

### Oocysts produced by F-RBP deficient females are sterile

We next sought to understand at which point loss of F-RBP halts the infection. Mice infected with F-RBP cKO parasites for six days were treated with rapamycin or vehicle for 72 hours, at which point fecal matter was collected, mice were sacrificed, and intestinal tissue was collected for punch biopsies and histology from the ileum of each animal (Figure 4A). PCR using parasite genomic DNA extracted from the intestinal tissue as template, demonstrated that at this time point the *f-rbp* locus remained intact upon vehicle treatment, but was fully excised following rapamycin treatment (Figure 4B). We had assumed that the serious fitness defect, and lower parasite burden in the intestine of rapamycin treated animals (Figure 4C) infected with the mutant was due to its inability to generate oocysts and thus measured the number of oocysts and their proportion of the overall parasite population using suitable antibodies for both by immunofluorescence. As expected, we detected much lower numbers of parasites in mice treated with rapamycin (Figure 4D). Surprisingly however, in relative terms oocysts appeared overrepresented in the diminished parasite population upon rapamycin treatment (Figure 4E). We thus wondered whether ablation of *f-rbp*, while allowing for oocyst production, renders those oocysts unable to initiate the next round of infection. To test this, we purified oocysts from the feces of groups of rapamycin treated and control animals. We readily detected oocysts in the feces of both groups (albeit in lower numbers for the treated animals) and importantly documented robust ablation of the locus by PCR (Figure 4F). *f-rbp* and `Δ`*f-rbp* oocysts were used to infect HCT-8 cell cultures, and nanoluciferase measurements were taken at 4h and 48h post infection to calculate parasite growth. *f-rbp* parasites showed growth but *`Δ`f-rbp* parasites did not (Figure 4G), and we conclude that loss of F-RBP leads to the production of sterile oocysts.

### Δ*f-rbp* sporozoites are unable to invade host cells

We sought to determine the basis of oocyst sterility and first considered that *`Δ`f-rbp* sporozoites may be unable to emerge from the oocyst. To initiate excystation, oocysts from both groups were resuspended in sodium taurodeoxycholate and sodium bicarbonate for 1h (27) and excystation was scored. We found that both *f-rbp* and *`Δ`f-rbp* were able to excyst with similar efficiency (Figure 4H). We then assessed the ability of *`Δ`f-rbp* sporozoites to invade host cells. Oocysts from both groups were excysted and allowed to interact with HCT8 cells for 2 hours, at which point cells were fixed and scored for successful invasion using a protection assay(28). We measured robust invasion for the vehicle treated parasites, that was sensitive to the known inhibitor latrunculin(29) (Figure 4I). In contrast, *`Δ`f-rbp* sporozoites showed a profound invasion defect. Interestingly, we observed even fewer intracellular parasites for the mutant than under latrunculin treatment, and overall, we noted very few parasites following *`Δ`f-rbp* exposure (Figure 4I). Latrunculin prevents actin polymerization, treated parasites attach but then fail to glide and invade. We therefore next measured parasite attachment and found *`Δ`f-rbp* sporozoites unable to attach to the host cells. Note that *f-rbp* sporozoites readily attach to their target, and that is not affected by Latrunculin treatment as previously shown(29) (Figure 4J).

We wondered whether these functional defects of the sporozoite are reflected in their morphology and mounted *f-rbp* and *`Δ`f-rbp* sporozoites onto poly-D-lysine slides, followed by staining with Hoechst and VVL. When observed by fluorescence microscopy and DIC we noted that *`Δ`f-rbp* parasites were significantly smaller than *f-rbp* sporozoites (p<0.0001, Figure 4K). By expansion microscopy *`Δ`f-rbp* sporozoites again appeared smaller and stouter with their apical structure less defined than their *f-rbp* counterparts (Figure 4L). We conclude that loss of F-RBP results in the shedding of sterile oocysts that carry defective sporozoites that are unable to attach to and invade epithelial cells.

### F-RBP binds a subset of transcripts highly expressed in females

F-RBP is predicted to act as an RNA binding protein based on the presence of a KH domain and RNA Recognition Motif(30). To test whether F-RBP indeed binds RNAs and to identify its targets, we performed RNA immunoprecipitation sequencing (31) using F-RBP-HA parasites. Western blot analysis of lysate from this strain subjected to anti-HA affinity matrix precipitation revealed a band of appropriate size (90 kDa). This band is absent when wild type lysate (WT) is used as control (Figure 5A, IP fraction). HCT-8 cell cultures were infected with F-RBP-HA parasites or control WT parasites and harvested after 54 hours and processed for immunoprecipitation and RNA sequencing. Reads were aligned to the *C. parvum* T2T genome(32) and using a statistical cutoff based on the log odds we determined that 57 parasite mRNAs and 2 lncRNAs were significantly enriched in the F-RBP immunoprecipitated samples (Figure 5B, Supplemental Figure 5A, Supplemental Table 2).

**Figure 5:**
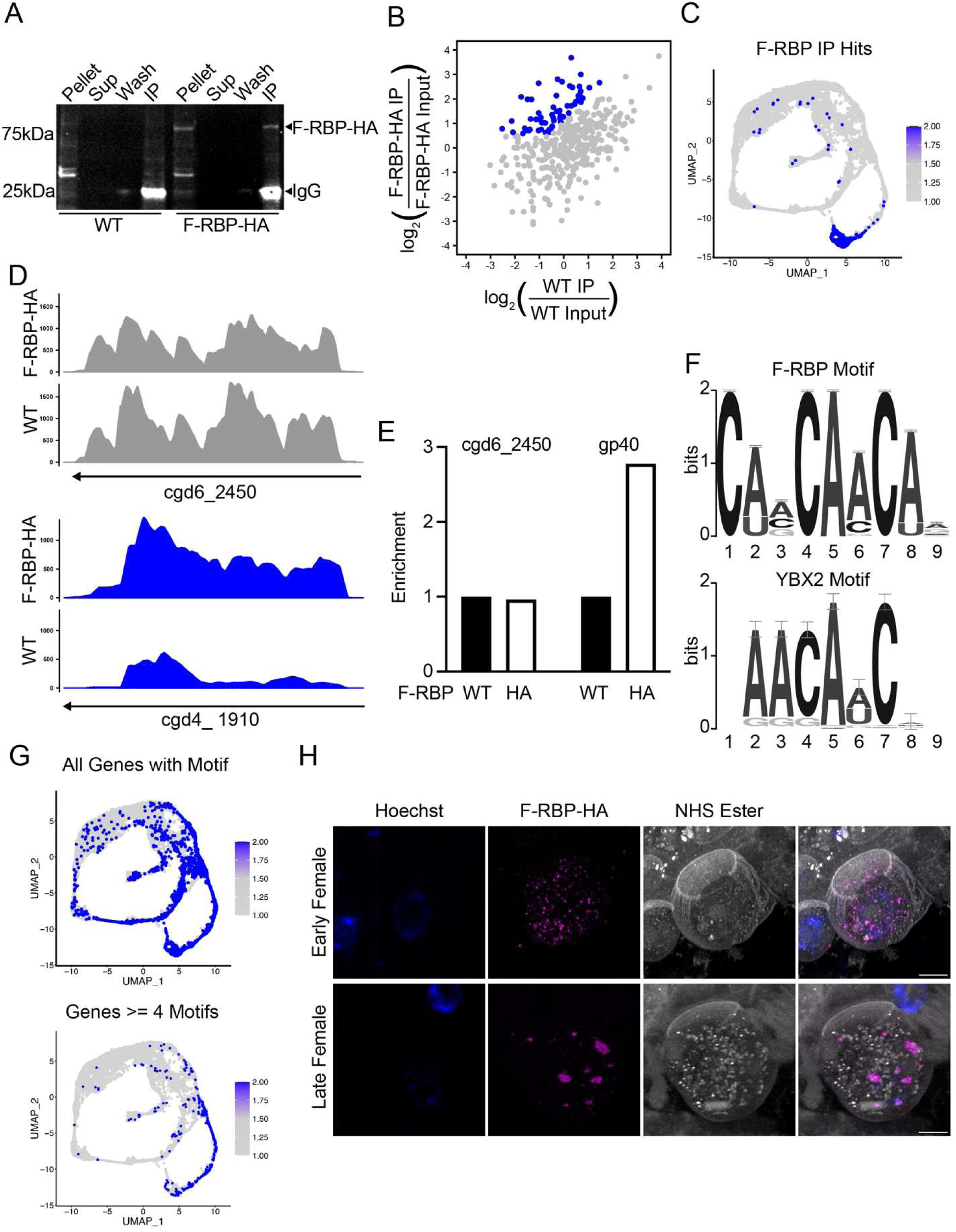
F-RBP binds specific transcripts upregulated in the female gamete and forms puncta in later females. **A.** Western blot detecting immunoprecipitation of F-RBP-HA from infected HCT-8 cells and wild type control using anti-HA antibody after 54 hours of infection. Lane annotations: Pellet (lanes 1 & 5) = cell pellet after sonication, Sup (lanes 2 & 6) = supernatant after IP, Wash (lanes 3 & 7)=washed affinity matrix prior to elution, IP (lanes 4 & 8) = immunoprecipitated sample. Note the bright band at 25 kDa in lanes 1 and 8 represent IgG from light chain of the HA antibody on affinity matrix. **B.** Relative enrichment of transcripts in the F-RBP and WT samples. Relative enrichment defined as the log_2_ of the ratio of IP and Input transcripts per million (TPM). Blue dots represent enriched transcripts in F-RBP-IP compared to the WT-IP. Non-enriched transcripts are represented in gray. **C.** The 57 protein coding RIP-Seq enriched transcripts plotted on the Single Cell Atlas as a module. Details on modfule score completion can be found in the Methods. **D.** Representative coverage plots for cgd6_2450 (a non- enriched transcript, top) and cgd4_1910 (an enriched transcript, bottom). Shown in counts per million. IP sample coverage plots for both shown. Gene model and direction for each gene represented as an arrow below the plot. **E.** RT-qPCR measurements of a non-enriched transcript (cgd6_2450) and an enriched transcript (cgd6_1080, gp40). Measurements were taken from a biological replicate. Enrichment was calculated using the percent input method. WT samples were normalized to an enrichment of 1. **F.** The MEME Suite logos representing the F-RBP motif (top) and most similar motif as determined by TOMTOM, YBX2 (bottom). **G.** All *Cryptosporidium* genes harboring the F-RBP motif (CAMCAACAR) (top) and genes with ≥4 copies (bottom) of the F-RBP motif plotted as modules on the Single Cell Atlas (top). **H.** U-ExM of female gametes performed on intestinal tissue sections. The earlier female (top) is identified by its smaller size and fewer wall forming bodies compared to the later female (bottom). Expansion factor 4.16. Scale bar 1µm.

Plotting the 57 protein-coding transcripts onto the *Cryptosporidium* Single Cell Atlas(6), showed enrichment for genes highly expressed late in the female gamete (Figure 5C & Supplemental Table 2) with 35 encoding proteins present in the sporozoite (Supplemental Table 2)(12). Not all genes highly expressed in females were enriched in the IP. For example, cgd6_2450 shows high transcript abundance in females, but is not enriched in the F-RBP IP (Figure 5D) or by RT-qPCR in a biological replicate (Figure 5E). However, the transcript of cgd4_1910 shows increased coverage in the F-RBP IP compared to WT IP (Figure 5D). Transcripts like cgd6_1080, encoding gp40, that show enrichment by RIPseq also showed enrichment when RT-qPCR was used to probe pull downs (Figure 5E). We conclude that F-RBP binds a subset of mRNAs expressed in female gametes.

### F-RBP targets share a sequence motif with targets of Y-Box family proteins

We searched for shared primary sequence features among the significantly enriched transcripts using the MEME suite(33). This resulted in the detection of a CAMCAACAR motif in the coding sequence of 27/57 of the enriched mRNAs (Figure 5F & Supplemental Table 3). A search for similar previously identified eukaryotic motifs using the TOMTOM tool(34) returned 21 matches with motif RNCMPT00084 standing out with highest significance (E-value and q-value <0.05). RNCMPT00084 corresponds to the YBX2 RNA-binding protein motif (E-value and q-value <0.05, Figure 5F, Supplemental Figure 6A, Supplemental Table 3)(35). YBX2 is a member of the Y-Box family of nucleic acid-binding proteins involved in a variety of developmental processes, but YBX2 is most notably associated with the development of gametes and early embryos(36, 37). Murine YBX2 is one of the most abundant maternal RNA binding proteins (37) and a crucial mediator of oocyte RNA stability and translation(36, 38). An unbiased scan of the *C. parvum* genome for the CAMCAACAR motif found that 937 protein coding transcripts contain at least one copy of this motif (Supplemental Table 4). These genes appear more highly expressed at the end of merogony and female gamete development and sporogony (Figure 5G). Among those that harbor multiple copies of the CAMCAACAR motif female transcripts are enriched and this enrichment is significant for the late female clusters 15-18 (Figure 5G & Supplemental Figure 7A for more detail). Y-Box proteins often aggregate into messenger ribonucleoprotein particles, RNA-protein complexes associated with mRNA stability and translational control(36). When we examined F-RBP localization at high resolution by expansion microscopy, we found a consistent developmental pattern. Early in female gametogenesis when the cells were small and harbored few wall forming bodies, F-RBP was diffusely distributed across the cytoplasm (Figure 5H). However, in more mature female gametes, identified by an abundance of wall forming bodies and larger cell size, F-RBP was aggregated into prominent cytoplasmic bodies (Figure 5H). We conclude that F-RBP interacts with mRNA, its targets are enriched for those carrying a conserved binding motif, and F-RBP form aggregates akin to bodies described in other eukaryotes.

## Discussion

Infection with *Cryptosporidium* is a major cause of severe diarrheal disease around the world(39–41). The infection contributes to early childhood mortality, but even less severe cases can result in malnutrition, stunting, and developmental delays(42–44). There is only one Food and Drug Administration-approved drug that lacks efficacy in those who need it most, immunosuppressed individuals and small children, and there are no vaccines available to protect children from the infection(45). The sexual phase of the parasite lifecycle has emerged has a promising future target for intervention, as disruption of parasite sex rapidly halts infection (6, 11) which findings reported in this study corroborate (Figure 3G, Supplemental Figure 4A). Mechanistic understanding of parasite sex is required to identify specific molecular targets for such intervention(5).

Gamete development and fate determination is a complex and highly dynamic process in *Cryptosporidium,* a single celled organism that lacks genetic sex determination in the form of mating types or sex chromosomes. Both male and female gametes are derived from the same asexual precursor cell with hard-wired timing(18). Evidence for circadian rhythm and the potential of molecular clocks in protozoan parasites has been growing recently (reviewed in(46)), and along with cell cycle count, and accumulation or depletion of a threshold factor, time is one of the conceivable drivers of transition to gamete development. In the related apicomplexan parasite *Plasmodium*, the transcription factor AP2-G controls the initial commitment to sexual fate (20, 21) and the subsequent differentiation into male and female gametes involves both protein factors and non-coding RNAs(47, 48). In *Cryptosporidium*, a commitment factor akin to AP2-G is not apparent, but the transcription factor Myb-M is necessary and sufficient to initiate the male program(6). A reciprocal factor for the female gamete has not been discovered, and female thus might be the default sex in the absence of Myb-M. Here we identified the *f-rbp* gene among the earliest female transcripts. However, time lapse imaging using a fluorescent reporter driven by the promoter of this gene demonstrated that it takes 5.5 h for the gene to be expressed, which represents almost the mid-point of the typical intracellular development cycle of this parasite. While fate commitment appears established when the merozoite leaves the last host cell(18, 49, 50), that fate is thus not immediately transcriptionally executed in the next host cell, but takes hours to manifest (Figure 6A). In broader terms, this appears the case for all stages of *Cryptosporidium*, where the first half of each intracellular cycle is occupied by shared housekeeping tasks, including ribosome biogenesis and the establishment of an interface suitable to export parasite effector proteins and import host metabolites (reviewed in(51)). Only when these tasks are completed, three distinct transcriptional programs emerge from the shared origin: asexual merogony, and male or female gametogenesis. The sex of the cell thus appears not as a primary overarching identity but manifests as alternative specialization halfway towards completion of intracellular development. We do not understand what defines fate prior to this point, or what initiates the transcriptional transitions following it. In *Plasmodium*, a long non- coding RNA serves as a negative regulator of sexual commitment(52, 53). *Cryptosporidium* transcribes numerous long non-coding RNAs with many showing stage-specific patterns of expression and they may have yet to be discovered regulatory functions(6, 32, 54). Epigenetic regulation of promoter accessibility through dynamic histone modification offers another model, one that is important to the regulation of lifecycle progression in other apicomplexans(55, 56).

**Figure 6:**
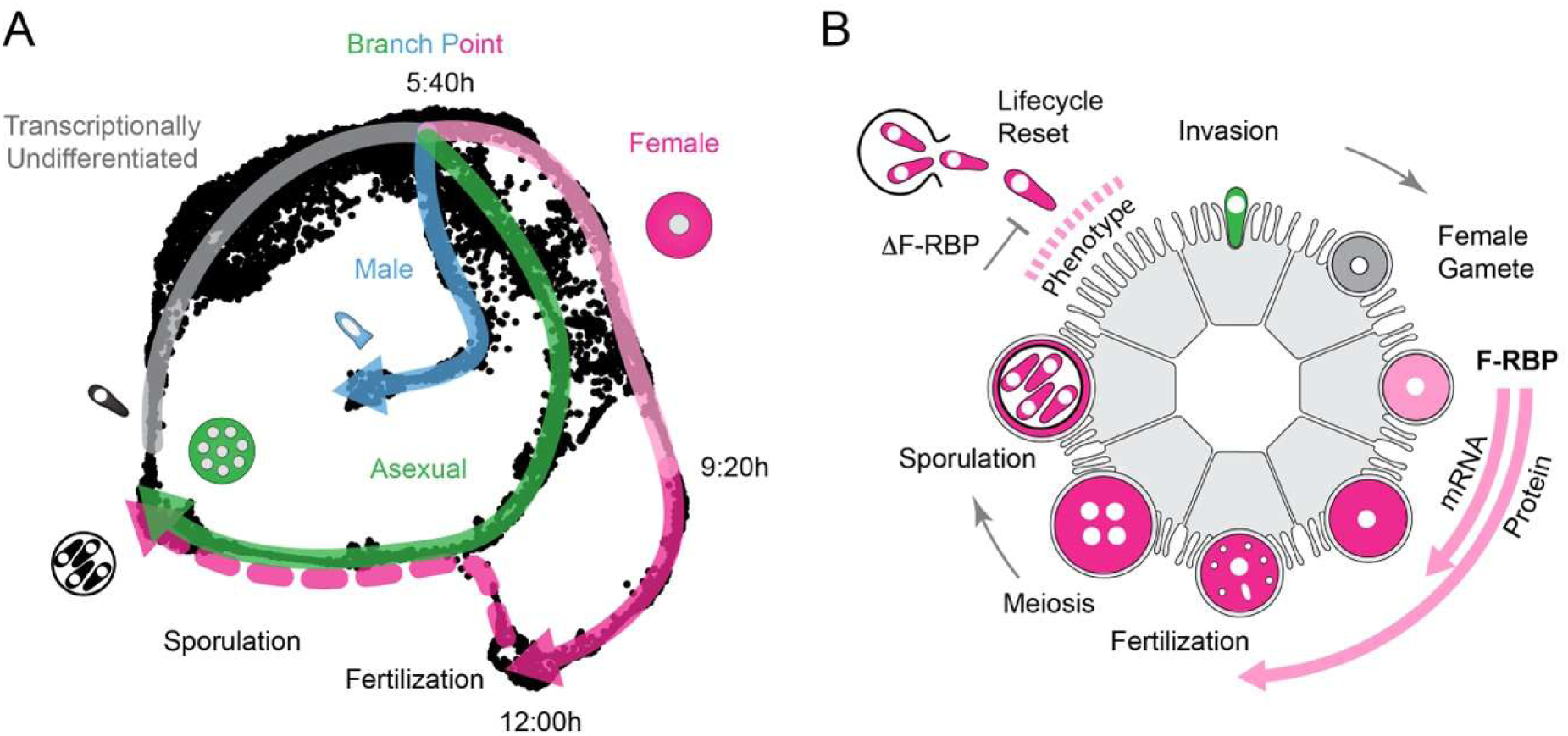
Transcriptional stage identity and F-RBP expression. **A.** Lifecycle model overlayed onto the single cell transcriptomic atlas(6). “Real” time indicated for female development based on the current study. Note an extended initial period for which the stage is not transcriptionally manifest. Following a branch point the expression of numerous stage specific genes leads to differentiated merozoites and gametes. Late assembly of sporozoites (dashed pink line) resembles that of merozoites. **B.** Schematic model of female development highlighting the presence of F-RBP mRNA and protein and the timing of the loss of function phenotype.

While initiation of transcription is a pivotal point to regulate gene expression in apicomplexans, post-transcriptional events are also of critical importance. Multiple RNA binding proteins were found critical to this process in *Plasmodium* and *Toxoplasma* (reviewed in(57–59)). Both rely on the Alba (acetylation lowers binding affinity) family of DNA/RNA binding proteins to regulate developmental processes(60). Genome mining failed to identify Alba homologs in *Cryptosporidium*. Both *Plasmodium* and *Toxoplasma* express multiple proteins that contain RRM and KH domains, the two domains also found in F-RBP(30, 61).

RNA binding proteins regulate the long-term storage and accurately timed sequential translation of mRNA that is crucial to the development of gametes and embryos across the eukaryotic tree(1, 62). Numerous such proteins are active across gamete development and embryogenesis, and several have been studied in *Plasmodium* (reviewed in(58)), often playing key roles in gamete development and fertility(47, 63). F-RBP is expressed very early in female development, but its loss does not cause defects in the development of female gametes (Figure 6B). Fate choice, fertilization and initial sporogony appear intact in Δ*f-rbp* mutants, however, the resulting oocysts are sterile (Figure 4). This suggests that F-RBP influences development after fertilization.

Several of the female RNA binding proteins studied in *Plasmodium* similarly function downstream of sexual differentiation and in the development of the zygote. Among the most intensely studied in this group is the RNA helicase DOZI (development of zygote inhibited) which, in complex with its binding partner CITH (car-I and trailer hitch), represses the translation of transcripts in mature female gametocytes whose protein products are needed later for zygote formation and meiosis(3, 64). Also in this complex is 7-Helix-1, and mutants lacking this factor show impaired gamete formation and reduced transmission of malaria parasites to mosquitoes(65). Puf2, another complex member participates in translational repression. Several Puf2 controlled transcripts encode ookinete surface proteins(66), and in *P. yoelii*, Puf2 is important while the sporozoite awaits transmission in the salivary gland and required for sporozoite infectivity (67). Ortholog and sequence similarity searches in *Cryptosporidium* reveal no homologs of Puf2 and 7-Helix-1. However, the *Cryptosporidium* genome does encode orthologs of DOZI and CITH (cgd8_1820 and cgd2_3220, respectively) and their transcripts appear present throughout most of the *Cryptosporidium* lifecycle.

Our study identified a *Cryptosporidium*-specific example of a maternal RNA binding protein required for successful development of mature sporozoites and transmission to the next host. More specifically, we showed that F-RBP forms cytosolic granules in later stage females, which was similarly seen for some of the proteins discussed above(3, 64, 65, 67). While many transcripts enriched in the immunoprecipitation of F-RBP encode for sporozoite proteins present in organelles such as the dense granules, micronemes and the crystalloid body a significant number remained unassigned by spatial proteomic analysis(12). Interestingly, sequence analysis of the mRNAs enriched by immunoprecipitation of F-RBP revealed a motif most similar to that of YBX2. YBX2 is a member of an ancient class of nucleic acid binding proteins found in both prokaryotes and eukaryotes (68). YBX2 in mammals is best known for regulating RNA homeostasis in gametes and early embryos and to assemble into cytoplasmic messenger ribonucleoprotein granules (36, 38, 69).

By what mechanism does maternal F-RBP ensure the production of infective sporozoites? F-RBP most likely regulates RNA homeostasis or translation. F-RBP may bind specific transcripts to influence their stability or decay. There are numerous examples of RNA binding proteins acting in this fashion in oocytes(2, 4). In *Plasmodium*, Puf2 has been implicated in mRNA decay (66) while DOZI is thought to act both in stabilization and decay of mRNAs(70). F-RBP may also act as a repressor of premature translation of female transcripts needed later for the development of sporozoites. This is a well-documented strategy of post-transcriptional control that egg cells use (1) and that was also found active in *Plasmodium* female gametocytes (reviewed in(59)) where mis-regulation of female translation was observed in the absence of Puf2 and DOZI(3, 66). mRNA homeostasis and translational control are not mutually exclusive mechanisms(66, 70) and it is likely that additional proteins are present alongside F-RBP in the granules we observe to aid in these tasks(3, 64, 65).

Ablation of F-RBP resulted in the first mutant in *Cryptosporidium* that results in a female sterile parasite. We demonstrate that successful control of gene expression over the complex lifecycle not only depends on timely transcription initiation facilitated by different transcription factors but also requires post-transcriptional processes that rely on stage specific RNA binding proteins and the formation of mRNA-protein granules.

## Materials and Methods

### Animals

Animal experiments were conducted with approval from the Institutional Animal Care and Use Committee of the University of Pennsylvania under protocol #8066292. *Ifny^-/-^* mice (RRID:IMSR_JAX:002287) were purchased from Jackson Laboratory and propagated as a breeding colony at the University of Pennsylvania. Moust housing conditions and age were as previously described(6).

### Plasmid construction

All primers used in this study can be found in Supplementary Table 1. Plasmids for genetic manipulation via CRISPR-Cas9 were generated as previously described(24, 71). Oligonucleotides (Sigma) encoding guide sequences were annealed and ligated into the *C. parvum* Cas9-U6 plasmid. Plasmids encoding the homology repair templates were made via Gibson assembly for the *f-rbp^mNeon^-cowp^tdTomato^*reporter strain, and restriction ligation cloning using AvrII and XmaI (New England Biolabs) for F-RBP-3xHA-eGFP, and BamHI and HindIII (New England Biolabs) for F-RBP-flox repair. Linear repair templates were generated by PCR using primers to encode recombination flanks of sequence identity to the target locus and PrimeSTAR Max DNA Polymerase Ver.2 (R047A, Takara).

### Generation of Transgenic Parasites

*C. parvum* IOWA II strain (IIa subtype) oocysts were purchased from Bunchgrass Farms and KVI parasites (IId subtype) (27)were maintained in-house through serial passage through *Ifng^-/-^* mice. Transgenic parasites were generated as previously described(24, 71, 72). Bunchgrass derived transgenic parasites were produced as described (6, 71). Transgenic parasites in the KVI background were produced as described(27). *Ifny^-/-^* mice were treated with antibiotics 4-7 days prior to infection and gavaged with 8% sodium bicarbonate immediately before oral gavage with transfected sporozoites. Selection for stable transgenics and oocyst shedding were performed as previously described(24). Oocysts were purified from feces by sucrose flotation and cesium chloride gradient and stored in PBS at 4°C(24, 71, 72). Oocyst genomic DNA was extracted with the Quick-DNA Fecal/Soil Microbe Miniprep Kit (Zymo) and integration of the repair template was validated by PCR.

### Cell line maintenance and preparation for infection

Human colorectal adenocarcinoma HCT-8 cells (ATCC, CCL-224TM) were maintained as previously described(6). Bunchgrass derived oocysts were prepared as described previously(71), resuspended in infection medium, and added to HCT-8 cells. KVI derived oocysts were incubated for 1 hour in 10mM HCl at 37°C and washed twice with PBS, then excysted onto host cells in media supplemented with 0.2 mM sodium taurodeoxycholate and 20mM sodium bicarbonate.

### Timelapse Fluorescence Microscopy

HCT-8 infections were set up in 8 well chamber slides (CellVis). Movies were acquired on a GE DeltaVision OMX (Penn Vet Imaging Core) at 5% CO_2_, and 37°C. Image stacks were acquired every 15 minutes for 16 hours. Movies were processed using GE softWoRx and Fiji. Fluorescence intensity was measured using Fiji for Mac OS.

### Immunofluorescence Assay (IFA)

For all HCT-8 infections: HCT-8 cells were grown in 96 well plates, washed, and fixed at time points indicated in the figure legends, and processed as previously described(6). Primary antibodies included rat anti-HA (1:1,000, Roche, 3F10), rabbit anti-HA (1:1,000, Cell Signaling Technology), VVL-Biotin (1:10,000, Vector Labs), VVL-fluorescein (1:1,000, Vector Labs, FL1231), mouse-anti DMC1 (1:10, a gift from C. Huston,(8)). For the protection assay, cells were treated with Latrunculin A (Abcam) at 2µM, 30 minutes prior to infection. Parasites were allowed to infect for 2 hours at 37°C prior to fixation. The staining protocol was adapted from(28). Cells were stained with 1:2000 VVL-Biotin (Vector Labs) in 3% BSA for 30 minutes, then washed 3 times for 3 minutes with dPBS. Samples were permeabilized with 0.01% Digitonin for 15 minutes and washed with PBS-0.01% Tween. For all HCT-8 experiments, Alexa Fluor secondary antibodies were used at a dilution of 1:1,000. Additional stains for the intracellular/extracellular and attachment assay include Sporo-Glo-A600FLR 1:20 in Sporo-Glo buffer for 1 hour followed by 1:1000 DAPI for 10 minutes. Coverslips were mounted with Fluorogel (Electron Microscopy Science).

For intestinal sections: Mice were sacrificed either at peak of infection or after designated treatment time. Intestinal tissue was “Swiss-rolled” and processed as previously described by the Penn Vet Comparative Pathology Core(73). Sections were stained the primary antibodies including rat anti-HA primary antibody (1:500, Roche, 3F10), Crypto-Glo-Biotin (1:20, Waterborne), rabbit anti-sporozoite polyclonal (1:1000,a gift from F. Tosini(74)). AlexaFluor secondary antibodies were used at a concentration of 1:250, with Hoescht at a concentration of 1:2000 Hoechst. Slides were mounted as described in the previous section.

For sporozoites: Parasites were excysted as described earlier and fixed in suspension with 4% PFA. Sporozoites were stained in suspension with VVL-fluorescein (1:2000,Vector Labs) and 1:500 Hoechst in PBS. Sporozoites were washed 2 times with PBS and transferred to an in 8 well chamber slide coated in poly-D Lysine (CellVis). Sporozoites were allowed to settle for 20 minutes, then washed with PBS and imaged.

For UEx-M: Intestinal samples were processed, sectioned, stored at -80°C until and expanded for U-ExM as described previously(75, 76). Primary antibodies include rat anti-HA 1:250 (1:250, Roche, 3F10). Secondary antibodies used were Alexa Fluor goat anti-rat 488 (1:500), NHS Ester Atto-647 (1:200, Sigma-Aldrich), and 1:1,000 Hoechst. For sporozoites: Parasites were excysted and were transferred to poly-D lysine coated coverslips and allowed to settle for 20 minutes. Sporozoites were spun at 200g for 1 minute, then coverslips were processed as described above. Dyes used were 1:1,000 Hoechst and NHS Ester Atto-647 (1:500, Sigma-Aldrich).

F-RBP-eGFP oocysts were imaged without fixation or stain after being allowed to settle on an Ibidi µ-Slide VI 0.4 (Thermo Fisher).

All images were taken on a Lecia DM6000 Widefield microscope (Penn Vet Imaging Core) using the 100x objective or a Leica Stellaris FALCON 31594 Confocal microscope (Penn Vet Imaging Core) using the 63x objective with Leica Application Suite X software. Images were processed and analyzed using Fiji for Mac OS software.

### Flow Cytometry

Oocysts were purified from feces using a miniaturized sucrose gradient and stained as previously described(25). Samples were processed on a LSRFortessa or FACSSymphony A3 Lite. Data were analyzed with FlowJo software (TreeStar).

### Generation of loss-of-function mutant

To generate the F-RBP flox plasmid, a recodonized gene fragment was synthesized (GenScript) including a multiple cloning site, upstream endogenously encoded region of homology to F-RBP, an intron containing a loxP site, the recodonized portion of F-RBP starting at base 1722 including the KH domain excluding the stop codon, and the downstream loxP site. This fragment was digested with BamHI and HindIII (New England Biolabs) and ligated into the *C. parvum* nanoluciferase-neomycin resistant plasmid(24). Homology repair template and guide RNA were produced as described(24, 71). Animals were co-infected with 5,000 oocysts each of parasites harboring the successful integration of this construct and parasites with the inducible Cre recombinase(25). Recombinant progeny were selected as previously described with paromomycin and BRD7929(6, 25). Oocysts were purified from feces collected from days 18-24 post-infection and propagated in *Ifny^-/-^* mice.

### Rapamycin Treatment

For *in vivo* experiments, mice were treated with 10 mg/kg rapamycin (Thermo Fisher Scientific) and equally diluted ethanol as previously described(6). Mice were treated daily by oral gavage from two days prior to infection to ten days after infection. For *in vitro* experiments, parasites were cultured in the presence of rapamycin or equally diluted ethanol as a vehicle control as previously described at the start of infection(6).

### RT-qPCR

For whole cell lysate samples: Infection was performed in a 24 well plate. HCT-8 cells were infected and treated with vehicle or rapamycin as described in a previous section. RNA was isolated using the Qiagen RNEasy Kit and cDNA was synthesized as previously described(6). cDNA was diluted 1:4 for all measurements. Reactions were conducted in SsoAdvanced Universal SYBER Supermix (Bio-Rad). The ΔΔ*C_t_* method (77) was used to determine relative expression with *C. parvum* GAPDH as the control gene. All qPCR primers have been previously published(78).

For RIP-seq validation: RNA Immunoprecipitation was performed as described in the RNA Immunoprecipitation and Sequencing section. RNA concentration was measured by qBit prior to cDNA synthesis. 48 ng RNA was used to generate cDNA. cDNA was produced as described in the previous section. The percent input method was used to determine enrichment between samples. Primers for gp40 (cgd6_1810) and cgd6_2450 were previously published(78).

RT-qPCR was performed using a ViiA 7 Real Time PCR System (ThermoFisher) with previously described conditions(6).

### Nanoluciferase Assay

Culture growth assay: In quintuplet, HCT-8 cells were infected and at 4h and/or 48 hours post-infection cells were lysed with lysis buffer (24) and mixed with 1:50 NanoGlo substrate (NanoGlo Luciferase Assay Kit, Promega) in an opaque white 96 well plate. Luminescence was measured with a Promega GloMax Plate Reader(24, 71). From feces: 20mg of feces was dissolved in 1mL of fecal lysis buffer and processed as previously described (24). From intestinal tissue: Ileum was removed, and three 5mm punch biopsies were taken from the most distal region of the ileum from each animal. Each sample was measured in triplicate, with final values calculated as the average of all samples for an individual animal.

### gDNA extraction and PCR of intestinal tissue

An intestinal section measuring approximately 1 cm in length was taken from each animal. The tissue was digested and DNA was extracted following the QIAmp DNA Mini Kit Qiagen. DNA was diluted 1:5 to perform PCR.

### Excystation Assay

50,000 oocysts were excysted as described earlier. The number of oocysts was determined immediately after triggering excystation using a hemocytometer (Kova Plastics). The number of resulting sporozoites was determined one hour after incubation at 37 °C. Percent excystation was calculated by dividing the sporozoite number by 4x the original oocyst count (assuming 4 sporozoites per oocyst).

### RNA Immunoprecipitation and Sequencing (RIP-Seq)

Infections were set up in duplicate in T25 flasks 24 hours prior to infection. In each flask, 4 x 10^6^ Bunchgrass or F-RBP-HA parasites were used. Media was changed to infection media and the flasks were incubated for 54 hours. Protocol for RNA Immunoprecipitation was modified from previous methods(79). Cells were fixed with 0.1% paraformaldehyde for 8 minutes at room temperature. The reaction was quenched by adding glycine to a final concentration of 125 mM for 5 minutes. Cells were removed from the flask and pelleted at 500g for 5 min 4°C. Cells washed twice with PBS and pelleted at 500g for 5 minutes. Cells were resuspended in RNP Lysis Buffer (20 mM Tris pH 7.5, 140 mM NaCl, 1.8 mM MgCl_2_, 0.1% IGEPAL, 0.1% SDS, 10% glycerol, 1mM DTT, 1x protease inhibitor cocktail EDTA free (Roche) and 2mM vanadyl ribonucleoside complex (NEB)). Cells were sonicated in three cycles of 10s on, 30s off at 20% power using a Sartorius Labsonic M sonicator. Sonicated samples were pelleted at 16,000g for 30 minutes at 4°C. Supernatant was removed at 10% of the volume was frozen at -80°C for the input samples. Vanadyl ribonucleoside complex was added to the remainder of the supernatant and was used for immunoprecipitation using an HA-affinity matrix (Roche). Samples were immunoprecipitated overnight and treated as described in the product manual. RNA was isolated by incubation in RNA Extraction Buffer (20 mM Tris-HCL pH 7.5, 5 mM EDTA, 50mM NaCl, 0.1% SDS and 50 ug mL^-1^ proteinase K) at 70°C for 40 minutes. Samples of the same genotype were combined to purify the RNA by phenol-chloroform extraction. Purified RNA was treated with DNase and purified again by phenol-chloroform extraction. This RNA was used to prepare cDNA-based, Illumina-compatible sequencing libraries using the Takara SMART-Seq Total RNA Pico Input with UMIs (ZapR Mammalian). The ribosomal cDNA depletion step of the SMART- Seq kit was omitted in order to minimize sample loss. Sequencing libraries were then converted into Ultima-compatible libraries using proprietary primers “UG-tR1 Indexing Primer v2, P1” and “UG-tR2 Universal Primer v2”, provided by Ultima Genomics. Sequencing was performed on the UG 100 platform by Ultima Genomics, generating between 130 to 160 million single-end reads per sample, with most reads ranging between 200-400 bp in length.

Reads were trimmed using cutadapt (80)(-u 22, -a NNNNNNNNNNNNNNAGATCGGAAGAG, -q 20, -m 50, -l 500) and aligned to the *C. parvum* Bunchgrass telomere-to-telomere genome (32) (T2T-CpBGF, alignment to mRNA and lncRNA loci only) using STAR(81). Aligned reads were further cleaned up using custom scripts that (a) remove ambiguous reads likely stemming from rRNA loci nested within coding-gene loci and (b) subset bam files for unique alignments only. The resulting cleaned-up unique alignments were used as input for salmon quant(82) (alignment- based mode, -l SR) to perform transcript quantification (transcript per million, TPM). Salmon output was then processed in a custom R script to identify putative F-RBP target transcripts. Only genes with more than five reads mapped in all four libraries were included for analysis. Using TPM values for all remaining genes, we computed the base-2 logarithm of TPM^IP^/TPM^input^ for both wildtype control as well as F-RBP-HA samples (see Figure 5B) as a proxy for enrichment of any given transcript in either pulldown. To identify putative F-RBP targets, we computed the base-2 logarithm of F-RBP-HA[TPM^IP^/TPM^input^]/WT[TPM^IP^/TPM^input^] and used the resulting F-RBP IP enrichment scores as input for mixture modeling using the R package mixtools(83) (normalmixEM, k = 2, maxit = 100000, arbvar = F), analogous to previous studies leveraging it for RIP-seq analysis(84, 85). Mixture modeling in this case assumes that the observed enrichment scores arise from a combination of two underlying distributions, here interpreted as putative F-RBP targets (high enrichment score) vs. all other *C. parvum* transcripts (lower enrichment score). The resulting gaussian curves are shown in Supplemental Figure 5. Mixtools assigns each transcript posterior probabilities of belonging to either underlying distribution. These are then used to calculate and odds ratio for each gene, allowing for model- based identification of putative F-RBP targets (LOD > 0). We further subset the resulting 86 candidates to only include genes with log_2_(F-RBP-HA[TPM^IP^/TPM^input^]) > 0.5 to produce a finalized list of 59 putative F-RBP targets / IP hits.

For RIP-seq coverage plots (Figure 5D), cleaned-up unique alignments were converted into CPM (counts per million)-normalized bigwig files using bamCoverage (deepTools, --normalizeUsing CPM, --binSize 1, --smoothLength 50). These bigwig files were then used as input in custom R scripts using packages Gviz, tracklater, GenomicRanges and IRanges to generate coverage plots.

For a detailed outline of RIP-seq data processing and analysis, refer to supplemental code provided on GitHub (https://github.com/amdaniels22/Cryptosporidium_F-RBP_RIPSeqandMotif)

### Motif analysis

The RIP-Seq gene list was filtered from the T2T-CpBGF(32) genome annotation using AGAT v1.7.0(86), and protein-coding genes were separated from lncRNA gene. Coding sequences (CDS), 5′ UTRs, 3′ UTRs, 150 bp upstream regions, 150 bp downstream regions, and protein sequences were extracted from mRNA annotations with AGAT suite. All sequence types were prepared for motif discovery analysis.

*De novo* motif discovery was performed using MEME suite v5.5.9 (33) with any number of repetitions per sequence (-mod anr). Separate runs were executed for DNA, RNA, and protein sequence types. Only the CDS RNA motif model produced statistically significant motifs under default p-value thresholds. Discovered motifs were compared using Tomtom from MEME suite.

The most significant motif model was scanned against all T2T-CpBGF’s protein coding genes’ CDS sequences using FIMO from MEME suite. The output was parsed to count motif occurrences per gene, and a master data matrix was generated using a custom Python script. This matrix integrated: (i) raw motif counts per CDS, (ii) gene lengths, (iii) motif counts normalized per 1 kb, (iv) mean FIMO p-values, q-values, and relative motif positions, and (v) presence/absence in the original RIP gene list. The resulting list of genes were stratified by motif counts per gene into four groups: 1 motif, 2 motifs, 3 motifs, and ≥4 motifs per CDS. Gene lists for each stratum were extracted from the master matrix using awk script. We focused our analysis on genes having ≥4 copies of this motif.

*C. parvum* single cell RNA seq data (GSE232438) were used for mapping motif enriched gene sets onto cells. A master module score was computed across all motif-containing genes (937 genes). Cells with master scores >0 were defined as the enriched cell set. Subgroup module scores were then computed for each motif-count stratum and masked to the enriched cell set (NA for non-enriched or subgroup-negative cells). UMAP feature plots were generated using Seurat v5 package(87).

To investigate the cellular populations enriched for genes containing the 4 or more copies of the discovered F-RBP motif, we performed cluster based analysis using the published single cell dataset(6). The absolute magnitude and proportion of ≥4 motif enriched cells per cluster were visualized using raw count and 100% stacked bar charts, respectively, with the proportion visualization used in Supplementary Figure 7. To statistically validate cluster specific enrichment, a one-tailed Fisher’s exact test was performed for each cluster comparing the number of enriched cells versus non enriched cells within that cluster against all other clusters combined. p-values were adjusted for multiple hypothesis testing using the Benjamini-Hochberg (BH) false discovery rate (FDR) method. Significance was defined as an adjusted p-value < 0.05.

## Data Availability

RIP-Seq data is available as an NCBI BioProject with the following project accession number: PRJNA1499133. Code used to process the RIP-Seq data and for motif analysis can be found on GitHub (https://github.com/amdaniels22/Cryptosporidium_F-RBP_RIPSeqandMotif).

## Acknowledgements

We thank Jessica Byerly, Eleanor Smith, Gracyn Buenconsejo, and Chole Tang for help with animal experimentation, Allison Cohen for ultrastructure expansion microscopy guidance, Gordon Ruthel and the Penn Vet Imaging Core for microscopy technical assistance, Daniel Cutillo for RIP-seq library preparation, Arunasalam Naguleswaran for guidance on the RIP-Seq protocol, and Ultima Genomics (Fremont,CA,USA) for providing free-of-charge sequencing services for the RIP-Seq experiment as part of their “Count on Us” initiative. This work was supported in part by grants from the National Institutes of Health to B.S. (R37-AI112427 and R01AI127798), Mark and Pamela Winter Infectious Disease Fellowship to A.M.D., Swiss National Science Foundation Fellowship P2BEP3_191774 and P500PB_211097 to S.S., and Damon Runyon Cancer Research Foundation (Merk Fellow, DRG 2491-23) to C.N.

---

**Supplemental Figure 1:**
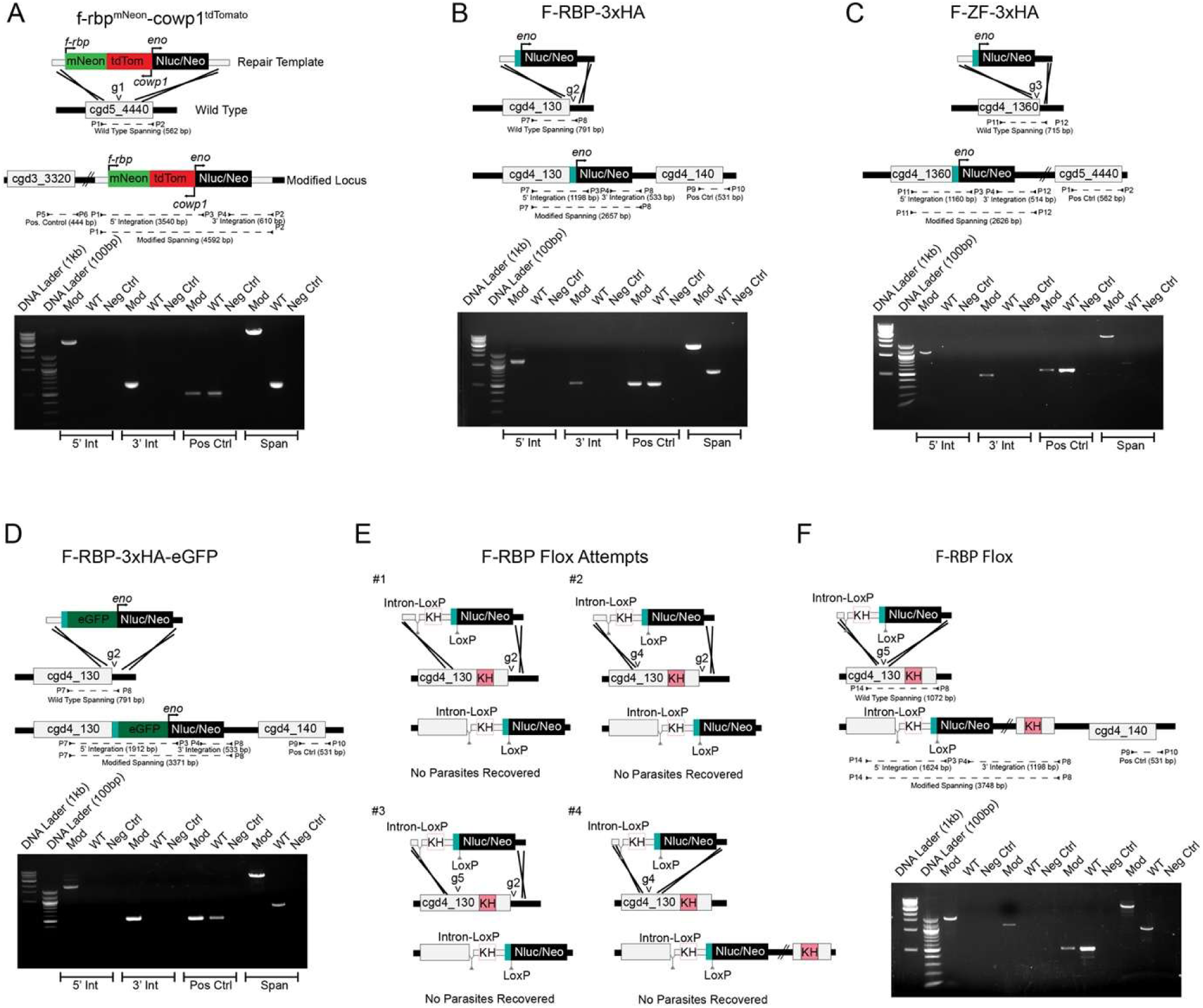
Generation of Transgenic Parasites. Schematics of native gene, homology repair template, and the modified loci. Area of the CRISPR-Cas9 induced break is denoted by arrowheads. PCR products showing desired integration are shown in the gene maps and on the gel, demonstrating correct homology repair. Strains include **A.** *f-rbp^mNeon^-cowp^tdTomato^.* **B.** F-RBP-3xHA. **C.** F-ZF-3xHA. **D.** F-RBP-3xHA-eGFP. **E.** F-RBP-flox unsuccessful attempts (no parasites recovered). **F.** F-RBP flox successful attempt. All strains were generated once except for F-ZF-3xHA which was generated twice. All primers and guides used can be found in Supplemental Table 1.

**Supplemental Figure 2:**
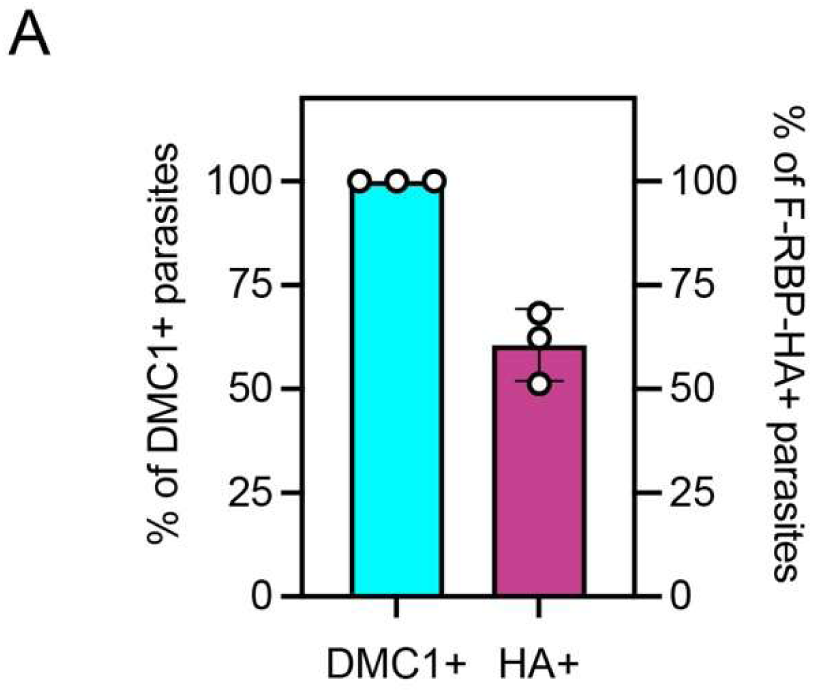
Differential timing of F-RBP-HA and DMC1 expression. **A.** Immunofluorescence quantification of the population of DMC1+ parasites expressing F-RBP-HA and F-RBP-HA+ parasites expressing DMC1 after 48 hours of HCT-8 infection with F-RBP-HA parasites. Note that all DMC1+ parasites expressed F-RBP-HA, but not all F-RBP-HA+ parasites were expressing DMC1. Representative of three independent experiments, where 3 coverslips were counted per experiment. 409 F-RBP-HA+ parasites counted across experiments.

**Supplemental Figure 3:**
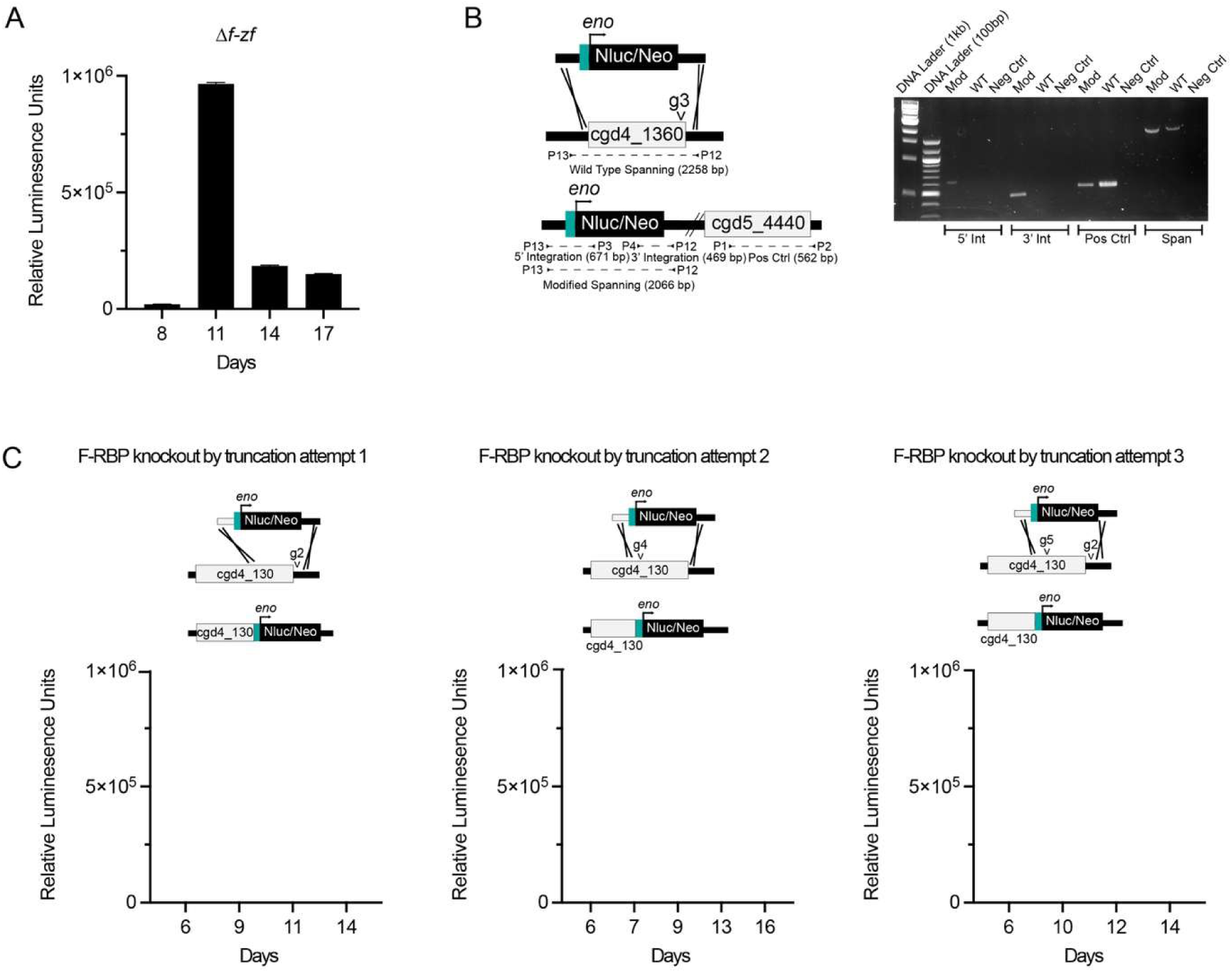
ΔF-ZF parasites were recovered, while F-RBP truncated parasites were not. **A.** Fecal luminescence measurements over time from the initial transfection of Δ*f-zf* parasites. **B.** Gene editing schematic for Δ*f-zf* parasites, including native locus, homology repair template, and modified locus (left) with corresponding diagnostic PCR (right). **C.** Fecal luminescence measurements for three separate transfection attempts at generating F-RBP truncated parasites with attempted truncation integration shown above each respective plot. Note these are separate attempt with different guide RNAs. All primers and guides used can be found in Supplemental Table 1.

**Supplemental Figure 4:**
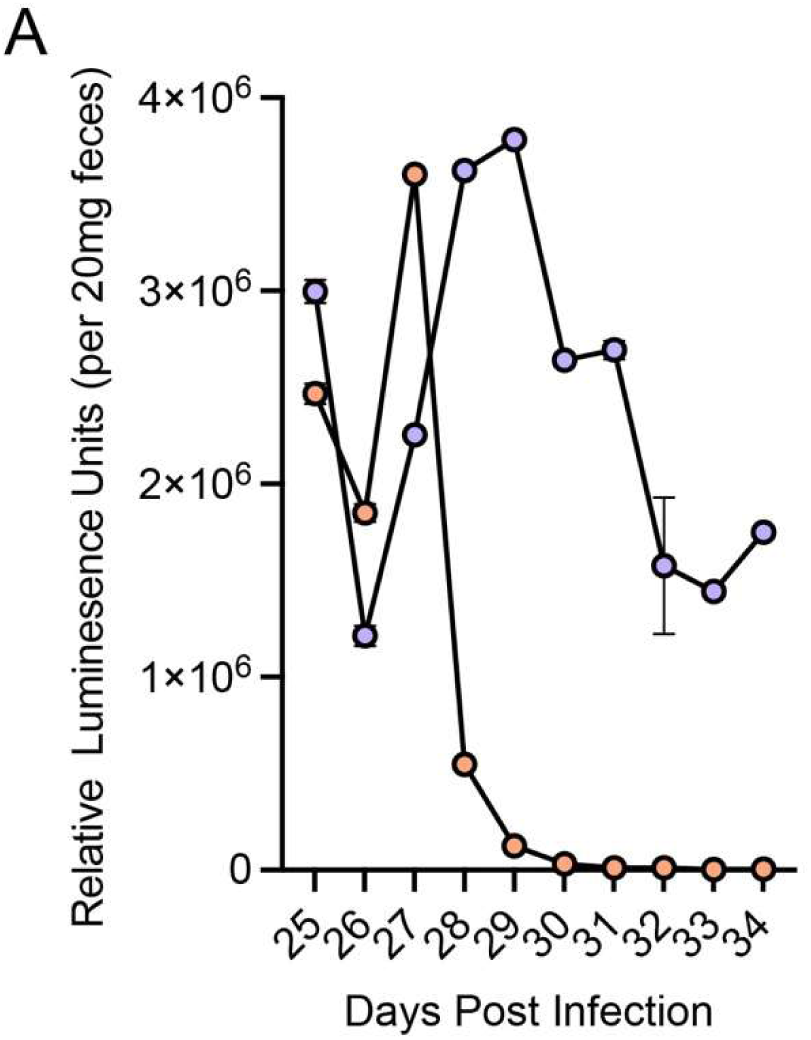
Rapamycin treatment of chronically infected F-RBP cKO mice cures animals from infection. **A.** Mice who were chronically infected with F-RBP cKO parasites were treated with vehicle (purple) or rapamycin (orange) were treated daily starting at day 25 post infection. Fecal luminescence was measured daily to assess oocyst shedding. 2-3 mice per treatment group.

**Supplemental Figure 5:**
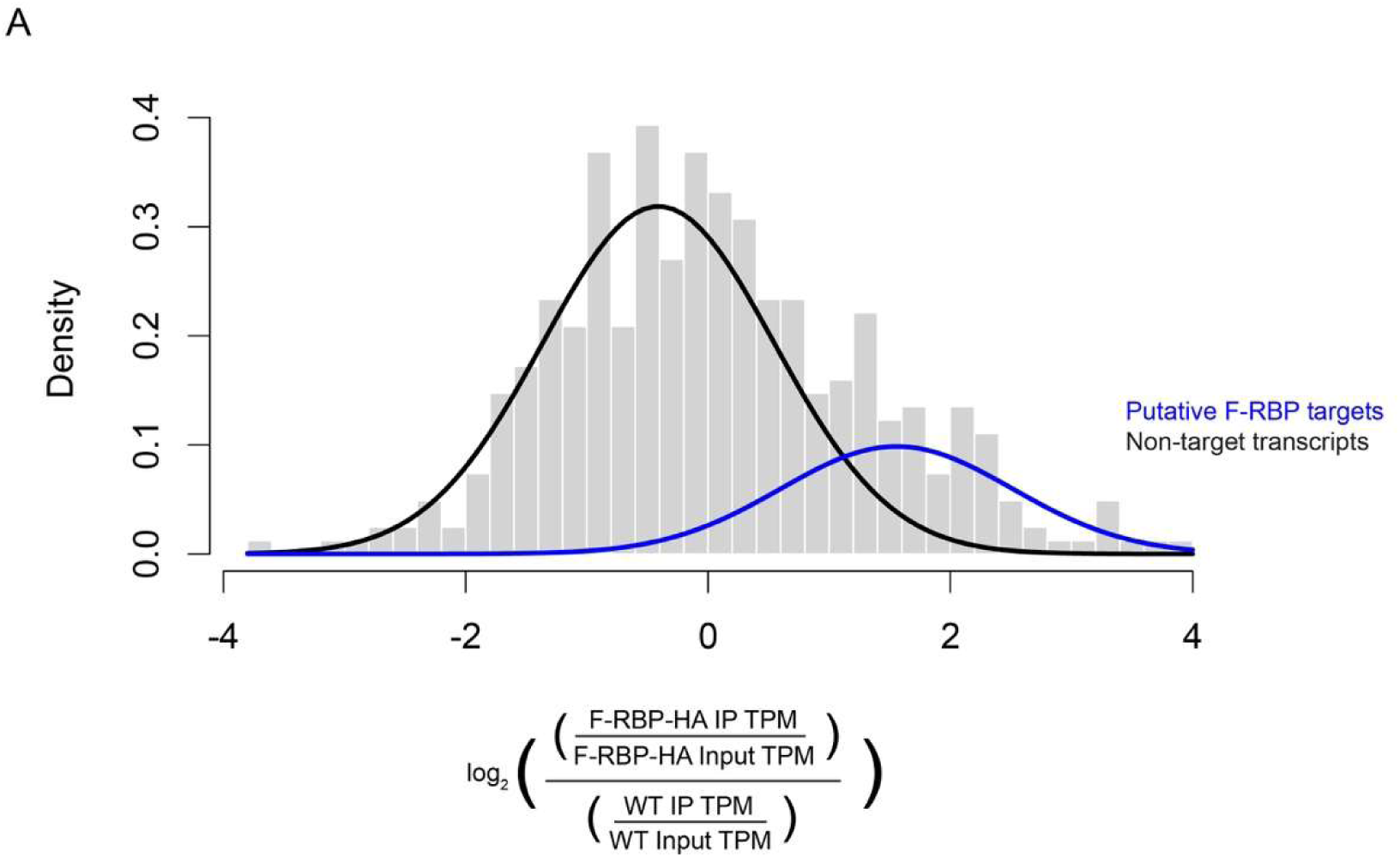
Gaussian mixture modeling for F-RBP RIP-Seq. **A.** Gaussian mixture model was calculated by utilizing F-RBP-HA IP enrichment scores based on the formula denoted on the X-axis. This modeling assumes that there are two distributions: F-RBP-HA IP enriched transcripts (blue) and other parasite transcripts (black). Details on model parameters and generation can be found in the methods and associated code.

**Supplemental Figure 6:**
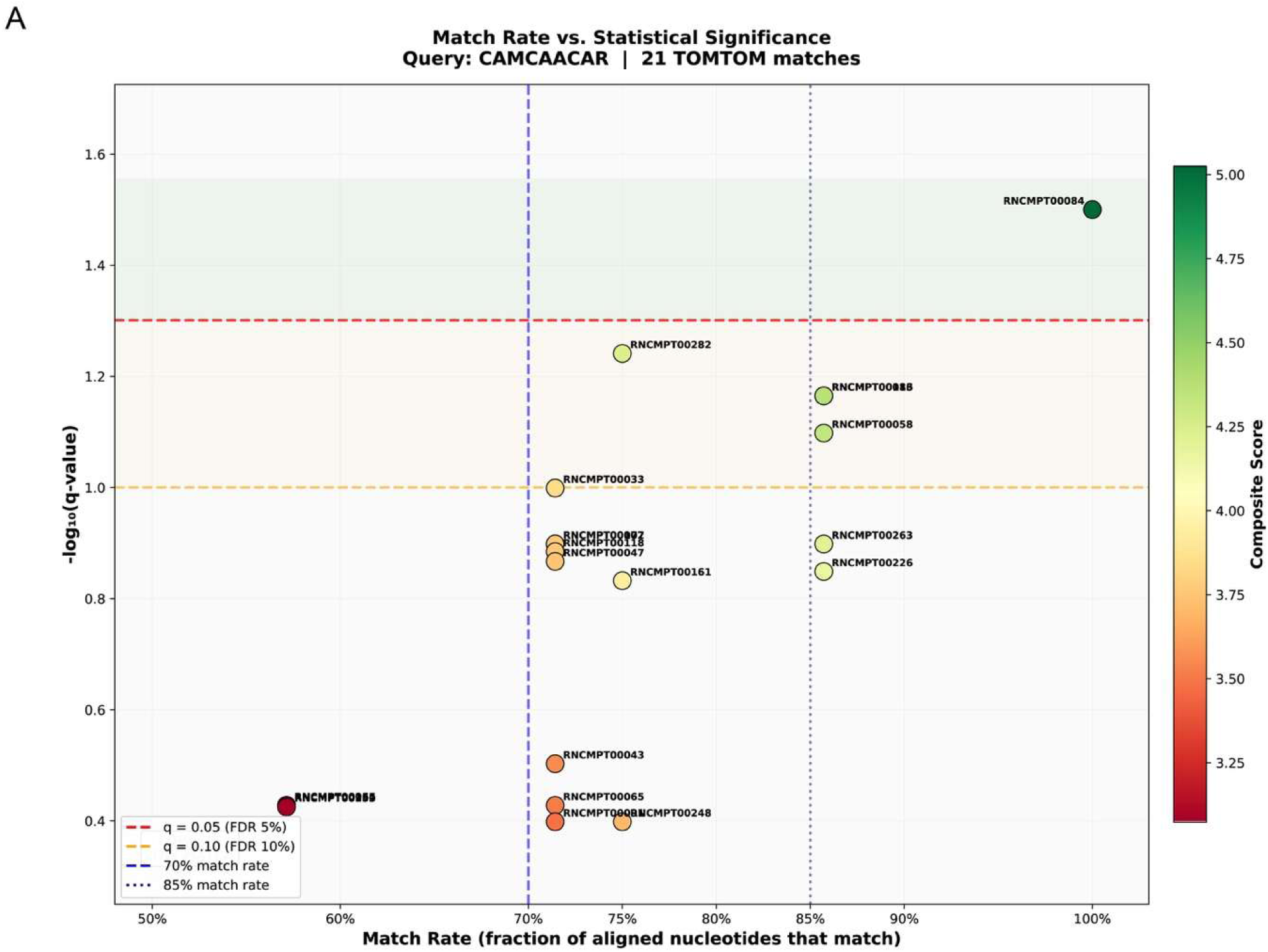
TOMTOM motif search returns only statistically significant similar motif, RNCMPT00084 (YBX2). **A.** 21 motif matches returned for TOMTOM search for CAMCAACAR similar motifs. Match composite score (scaled red to green) derived from a combination of q-value, E-value, p-value and match rate. -log_10_(q-value) and match rate shown in the graphic. All reported statistics can be found in Supplemental Table 3.

**Supplemental Figure 7:**
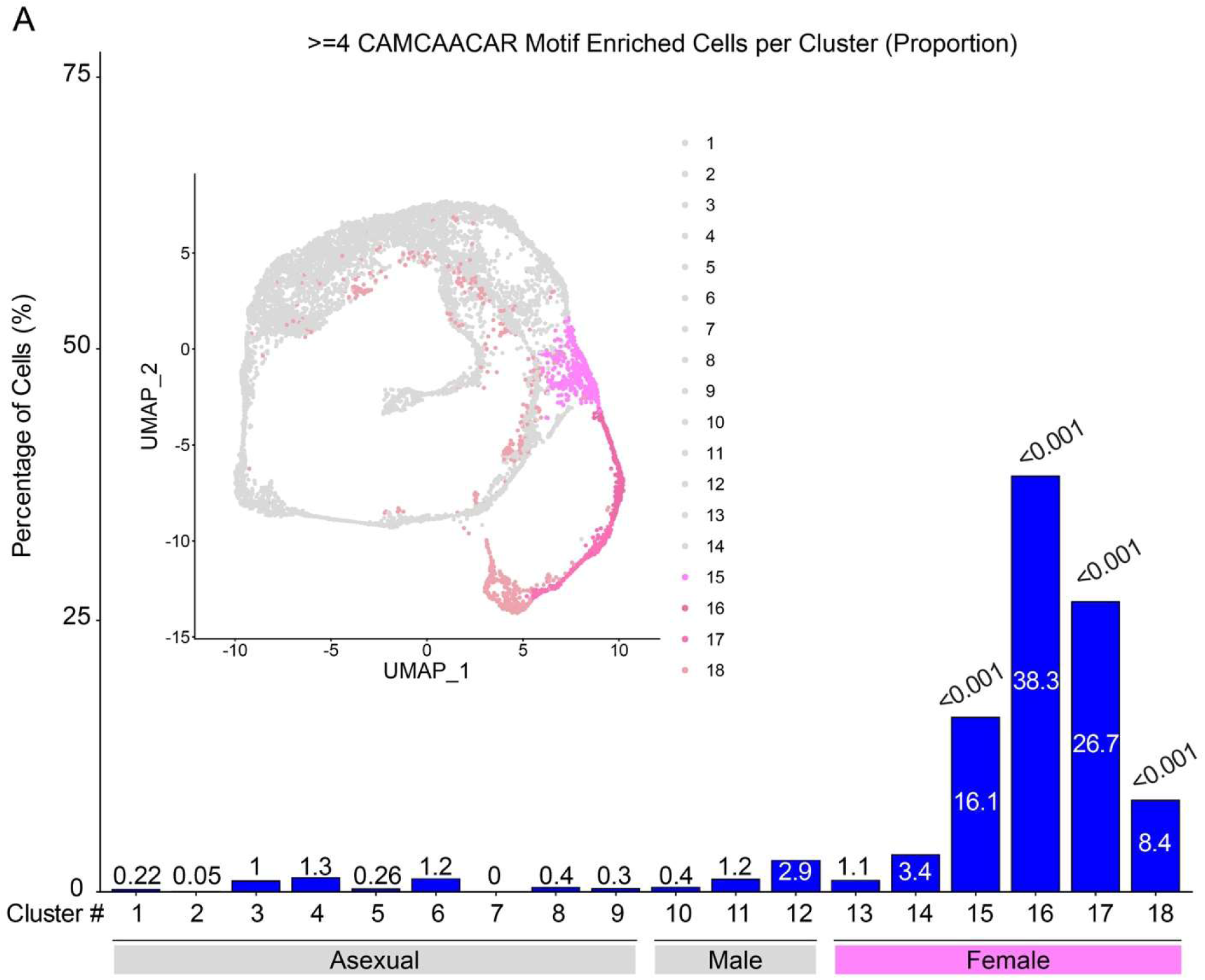
**Transcripts with ≥4 CAMCAACAR motifs are enriched in female gametes**. **A.** Percentage of cells (blue) which are enriched for genes containing ≥4 CAMCAACAR motifs separated by cluster of the *Cryptosporidium* Single Cell Atlas(6). Female clusters are denoted in pink on the inset UMAP projection and x axis, all other clusters represented in gray. Clusters 15,16,17, and 18 display significant enrichment. Adjusted P-values shown from a Benjamini-Hochberg False Discovery Rate analysis. All genes returned from the unbiased scan of the *C. parvum* genome with the CAMCAACAR motif and the statistics associated with the ≥4 motifs per transcript analysis can be found in Supplemental Table 4.

